# Conserved TIR-only proteins drive transcriptional defense and basal immunity in dicot and monocot plants

**DOI:** 10.64898/2026.06.07.730676

**Authors:** Henriette Laessle, Oliver Johanndrees, Jingqi Chen, Sabine Haigis, Tak Lee, Yu Chen, Li Liu, Wen Song, Jaqueline Bautor, Jan Jirschitzka, Bruno Huettel, Li Wan, Federica Locci, Jane E. Parker

## Abstract

Toll/interleukin-1/resistance (TIR) domain NADase enzymes signal in plant immunity by producing ribosylated nucleotide second messengers which activate EDS1 dimer-helper NLR pathways to restrict pathogen growth. Members of a small, distinctive group of TIR-only proteins are conserved between dicots and monocots, yet their functions remain poorly understood. Here, we show that conserved TIR-only proteins in Arabidopsis and barley share a fundamental enzymatic role in promoting basal defense against virulent filamentous pathogens, independently of NLR mediated effector-triggered immunity. Metabolite analysis of transiently expressed Arabidopsis and barley TIR-only proteins revealed their capacity to produce ribosylated cyclic nucleotides *in vivo*. By comparing phenotypes of *tir-only* and *eds1 pad4* CRISPR mutants in the two species, as well as *adr1* mutants in barley, we established that the TIR-only proteins promote PAMP-triggered transcriptional defenses associated with pathogen restriction. Barley possesses just one essential TIR-only enzyme and mutations of the two conserved *TIR-only* members in Arabidopsis were not compensated for by numerous other TIR-domain genes in the basal immune response. These findings suggest that conserved TIR-only proteins make a crucial contribution to TIR basal defense signaling networks of Arabidopsis and barley. We propose that a shared function in transcriptional defense regulation could explain the evolutionary retention of this discrete TIR-only group across monocot and dicot lineages.

## Introduction

The Toll/interleukin-1/resistance (TIR) domain has a central role in innate immunity signaling in plants, mammals and bacteria. In mammals, most TIR domains serve as self-associating adaptor modules in defense signal transduction (Nimma et al., 2017). A different immune activity of self-associating TIR domains as NAD^+^ hydrolyzing (NADase) enzymes was first discovered for mammalian sterile alpha and TIR motif containing 1 (SARM1) which promotes neuronal cell death through hydrolytic depletion of NAD^+^ (Essuman et al., 2017; Shi et al., 2022). Further bacterial and plant TIR-domain NADases have been characterized which hydrolyze NAD^+^ to produce cyclic and non-cyclic nucleotide immune-modulating signals (Ofir et al., 2021; Manik et al., 2022; Li et al., 2023; Locci et al., 2023).

A major family of intracellular nucleotide-binding/leucine-rich-repeat (NLR) immune receptors called TIR-NLRs (or TNLs) present in dicotyledonous (dicot) plants is defined by their N-terminal TIR signaling domain (Jones et al., 2016; Lapin et al., 2022). The TNLs, and members of a second large NLR receptor family possessing N-terminal coiled-coil (CC) signaling domains (CC-NLRs or CNLs), detect disease-promoting virulence factors (effectors) delivered by pathogens into host cells (Jones et al., 2016). NLR recognition normally blocks an effector from interfering with basal resistance responses mediated by cell-surface pattern-recognition receptors (PRRs) recognizing microbe- or damage-associated molecular patterns (MAMPs or DAMPs) (Couto and Zipfel, 2016; Jones et al., 2016). Effector-triggered immunity (ETI) is an amplified form of pattern-triggered immunity (PTI) and often results in host cell death at pathogen infection sites (Cui et al., 2015; Jones et al., 2016). Many pathogen effector-stimulated TNLs and CNLs assemble into signaling-active oligomers called resistosomes (Contreras et al., 2022; Chai et al., 2023; Hu and Chai, 2023; Guo et al., 2026). Studied TNLs form a tetrameric resistosome in which two asymmetrically arranged TIR domain pairs create a composite nicotinamide adenine dinucleotide hydrolase (NADase) catalytic site binding NAD^+^ and ATP substrates (Ma et al., 2020; Martin et al., 2020) to generate cyclic and non-cyclic ribosylated nucleotide intermediates (Horsefield et al., 2019; Wan et al., 2019; Huang et al., 2022; Jia et al., 2022; Bayless et al., 2023). By contrast, characterized CNL resistosomes of various oligomeric sizes can form calcium (Ca^2+^) permeable ion channels at host membranes which directly elevate cytoplasmic Ca^2+^ levels to promote host transcriptional defense and cell death (Hu and Chai, 2023; Guo et al., 2026). A number of TNL, CNL and CNL-like resistosomes, referred to as helper NLRs, function downstream of pathogen-sensing cell-surface PRRs and intracellular NLRs to amplify host defenses as part of a mutually reinforcing NLR-PRR immune signaling network to block infection (Feehan et al., 2020; Yuan et al., 2021; Jones et al., 2024; Locci and Parker, 2024).

Dicot plants also express numerous truncated TIR-nucleotide binding (TIR-NB) and TIR-only proteins with immune signaling roles (Meyers et al., 2002; Nandety et al., 2013; Lapin et al., 2022). Many have NADase-dependent cell death inducing properties in *N. benthamiana* transient expression assays (Nandety et al., 2013; Johanndrees et al., 2022; Bayless et al., 2023; Bayless et al., 2025). Arabidopsis TIR-only protein RECOGNITION OF HOPBA1 (*At*RBA1) confers resistance to a bacterial effector HopBA1 (Nishimura et al., 2017). Other Arabidopsis TIR-NB and TIR-only genes are rapidly upregulated in pattern-triggered basal immune responses (Tian et al., 2021), indicative of convergent roles of pathogen-induced TIRs in coordinating PTI and ETI (Ngou et al., 2021; Pruitt et al., 2021; Tian et al., 2021; Yuan et al., 2021).

TNL resistosomes, TIR-only proteins, and bacterial TIR-domain enzymes, employ the same fundamental mechanism requiring a conserved catalytic glutamic acid (Glu) residue to convert NAD^+^ and ATP substrates into nicotinamide adenine mononucleotide (NAM), ADP-ribose (ADPR), and cyclic ADP-ribose (cADPR) molecules, 2’cADPR and 3’cADPR (Horsefield et al., 2019; Wan et al., 2019; Ofir et al., 2021; Manik et al., 2022; Bayless et al., 2023). Two sets of plant TIR-catalyzed non-cyclic ribosylated nucleotides: ADP-ribosylated-ATP (ADPr-ATP)/ADPr-ADPR (di-ADPR) and 2’(5’’-phosphoribosyl)-5’-AMP/ADP (pRib-AMP/pRib-ADP), respectively, bind to and activate dimers of ENHANCED DISEASE SUSCEPTIBILITY 1 (EDS1) with SENESCENCE-ASSOCIATED GENE 101 (EDS1-SAG101) and EDS1 with PHYTOALEXIN DEFICIENT 4 (EDS1-PAD4) (Huang et al., 2022; Jia et al., 2022). The nucleotide-bound EDS1-SAG101 and EDS1-PAD4 complexes are then specifically recognized by N REQUIREMENT GENE 1 (NRG1) and ACTIVATED DISEASE RESISTANCE 1 (ADR1) CC-like helper NLRs, respectively (Huang et al., 2022; Jia et al., 2022; Yu et al., 2024; Huang et al., 2025; Xiao et al., 2025). Current evidence suggests that these activated helper NLRs also form membrane-associated resistosomes with Ca^2+^ ion channel activity to promote host transcriptional defense and cell death (Jacob et al., 2021; Wu et al., 2021; Wang et al., 2023; Yu et al., 2024; Huang et al., 2025; Wang et al., 2025; Xiao et al., 2025).

In dicot plants such as Arabidopsis and *N. benthamiana* (*Nb*), the ADPr-ATP/di-ADPR stimulated EDS1-SAG101-NRG1 (ESN) node promotes TNL mediated host cell death in ETI (Qi et al., 2018; Gantner et al., 2019; Lapin et al., 2019; Saile et al., 2020; Sun et al., 2021; Feehan et al., 2023). By contrast, the pRib-AMP/pRib-ADP stimulated EDS1-PAD4-ADR1 (EPA) node has a broader role in amplifying defense (without cell death) downstream of TNLs and certain CNL receptors (Lapin et al., 2020; Jacob et al., 2023; Wang et al., 2024; Ge et al., 2026), as well as cell-surface PRRs such as Arabidopsis RECEPTOR-LIKE PROTEIN 23 (RLP23) recognizing the oomycete MAMP necrosis-and ethylene-inducing peptide 1-like 20 (nlp20) (Pruitt et al., 2021; Tian et al., 2021).

Whereas dicot genomes contain numerous TIR-domain genes with diverse domain architectures, monocot genomes lack TNLs and TNs but retain a small number of conserved TIR-NB-tetratricopeptide-repeat (TNP) and TIR-only genes (Yue et al., 2012; Johanndrees et al., 2022; Liu et al., 2023). Phylogenetic and Hidden-Markov Modelling (HMM) studies placed the TIR-only group shared by dicots, monocots and magnoliids into a separate clade from other TIR-only, TN and TNL proteins (Johanndrees et al., 2022; Lapin et al., 2022). Transient over-expression assays of conserved TIR-only proteins from Arabidopsis, barley (*Hordeum vulgare*), rice (*Oryza sativa*), *Brachypodium distachyon* (*Bd*) and maize (*Zea mays*) in *Nb* induced cell death requiring the catalytic Glu residue (Wan et al., 2019; Johanndrees et al., 2022; Zhang et al., 2025), indicating that this class of dicot and monocot TIR-only proteins share NADase activity. Moreover, an immune signaling study in monocot rice revealed that pathogen resistance triggered by CNL Pish recognizing *Magnaporthe oryzae* (*Mo*) fungal rice blast strain YN2, as well as PRR-mediated defenses in response to chitin and flagellin (flg22) elicitors, employ a single conserved TIR-only protein (*Os*TIR) catalyzing pRib-ADP stimulation of the EPA node to promote immunity (Wu et al., 2024). *Os*TIR and *Nb*TIR proteins are repressed by Ca^2+^-binding sensors in healthy (pathogen non-triggered) tissues and released upon pathogen attack to activate EPA mediated pathogen restriction in rice and stomatal immunity in *Nb* (Wang et al., 2024; Wu et al., 2024). Despite these parallels, the extent to which this small group of conserved TIR-only proteins have analogous immune functions in monocots and dicots remains unresolved.

Here we perform a comparative analysis of the conserved TIR-only protein group in dicot Arabidopsis accession Col-0 (containing two members, AT1G52900 and AT1G61105) and monocot barley cultivar Golden Promise (GP, with one TIR-only member, HORVU.MOREX.r3.2HG0134190). We examine the immune phenotypes of CRISPR-Cas9 generated *tir-only* and *eds1-pad4* mutants in both species and measure the capacities of their TIR-only proteins to hydrolyze NAD^+^ in vivo. We establish that while conserved TIR-only proteins in Arabidopsis and barley are dispensable for ETI, they are essential drivers of PAMP-induced transcriptional defense and basal immunity protection against virulent filamentous pathogens. We propose that a shared fundamental role in immune potentiation underlies the persistence of this discrete TIR-only group in dicot and monocot lineages.

## Results

### Shared and distinctive *At*TIR and *Hv*TIR cell death inducing properties in *Nb*

Of the two Arabidopsis conserved TIR-only paralogs, *AT1G52900* (hereafter *At*TIR), but not *AT1G61105* (TAIR), is transcriptionally upregulated in immune-triggered tissues (Johanndrees et al., 2022). We therefore compared the enzymatic and immune signaling activities of *At*TIR with barley TIR-only protein *Hv*TIR (HORVU.MOREX.r3.2HG0134190; Ensembl Plants).

We first measured enzymatic contributions of yellow fluorescent protein (YFP)-tagged *At*TIR and *Hv*TIR transiently over-expressed by agroinfiltration in leaves of *Nb* WT or the immune-signaling deficient *Nb eds1 sag101a sag101b pad4* (*Nb epss*) mutant (Lapin et al., 2019), using YFP as a negative control. Both *At*TIR and *Hv*TIR induced cell death at 2 d post infiltration (dpi), with *Hv*TIR eliciting stronger ion leakage (measured by conductivity) and macroscopic cell death than *At*TIR in leaves (Figure 1A, B). To determine whether NADase activity is required for the *AtTIR* and *HvTIR* cell death in *Nb*, we substituted their respective catalytic Glu residues with alanine (producing *At*TIR^E122A^ and *Hv*TIR^E128A^). Neither TIR catalytic mutant elicited ion leakage above the YFP control (Figure 1A, B). In contrast to *At*TIR, *Hv*TIR cell death was not compromised in the *Nb epss* mutant (Figure 1A, B), indicating that transiently over-expressed *Hv*TIR elicits cell death independently of *EDS1* signaling. This could not be explained by higher *Hv*TIR protein accumulation than *At*TIR in these *Nb* assays (Supplementary Figure S1A), although it is possible that reduced *Hv*TIR protein levels result from enhanced cell death-associated turnover. We considered whether the *Hv*TIR cell death phenotype might be due to enzymatic depletion of NAD⁺ (Wan et al., 2019; Eastman et al., 2022; Bayless et al., 2023). To test whether this response was *Hv*TIR dosage dependent, we lowered the optical density (OD_600_) of *Agrobacteria* delivering *Hv*TIR to *Nb epss* leaves and found that this shifted cell death responses from *EDS1*-independent to *EDS1*-dependent (Supplementary Figure S1B). These data suggest that the *EDS1* requirement for *Hv*TIR induced cell death depends on the level of *Hv*TIR activity and that high catalytic output during transient overexpression in *Nb* may promote *EDS1* independent cell death, potentially through NAD^+^ depletion or the generation of additional, as yet unidentified, NAD^+^ derived metabolites.

**Figure 1:**
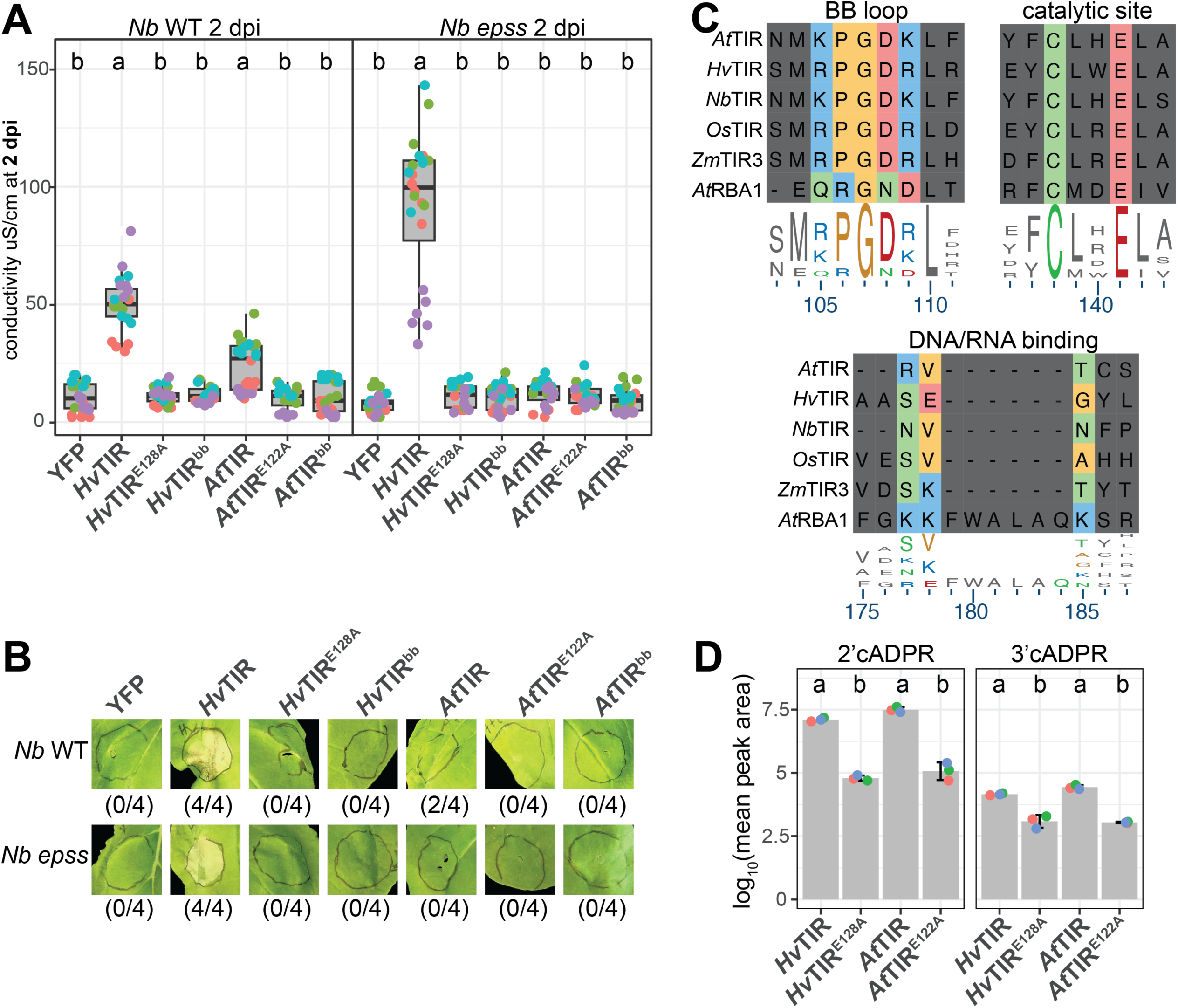
Overexpression of *Hv*TIR results in *EDS1*-independent cell death in *Nicotiana benthamiana*. (**A**) Ion leakage (conductivity in µS/cm) was measured as a proxy of cell death in 4–5-week-old *Nb* WT and *eds1 pad4 sag101a sag101b* (*epss*) plants. Leaves were infiltrated with *Agrobacteria* expressing *35S::HvTIR-YFP* (*Hv*TIR), *35S::HvTIR^E128A^-YFP* (*Hv*TIR^E128A^), *35S::HvTIR^bb^-YFP* (*Hv*TIR^bb^), *35S::AtTIR-YFP* (*At*TIR), *35S::AtTIR^E122A^-YFP* (*At*TIR^E122A^), *35S::AtTIR^bb^-YFP* (*At*TIR^bb^), and *35S::YFP* (YFP, as a negative control). Ion leakage was measured 2 d post infiltration (dpi). Constructs with a different letter code show statistically significant differences in conductivity (Kruskal -Wallis test with Nemenyi post-hoc, α = 0.05, with n = 24 from f our independent experiments). Individual data points are shown, with colors indicating independent experiments. For boxplots, the ce nter line indicates the median, edges show the 25th and the 75th percentiles, and whiskers indicate 1.5 × IQR. (**B**) Macroscopic cell death in *N. benthamiana* (*Nb*) WT and *Nb eds1 pad4 sag101a sag101b* (*Nb epss*) induced by *Agrobacterium*-mediated expression of *At*TIR and *Hv*TIR tagged with YFP and driven by the 35S promoter. Constructs include WT, non -catalytic (*At*TIR^E122A^, *Hv*TIR^E128A^), and BB -loop mutants ( *At*TIR^bb^, *Hv*TIR^bb^). Images at 3 dpi show representative cell death phenotypes. Numbers in parentheses indicate the number of independent experiments showing cell death out of four performed (one leaf per experiment). (**C)** Sequence alignment of conserved TIR-only protein motifs from Arabidopsis (*At*TIR), barley (*Hv*TIR), *N. benthamiana* (*Nb*TIR), rice (*Os*TIR), *Zea mays* (*Zm*TIR3) and Arabidopsis *At*RBA1 using Clustal Omega. Key amino acids highlighted. **(D)** Quantification via LC - MS of 2’cADPR and 3’cADPR in *Nb epss* leaves transiently expressing *Hv*TIR, *Hv*TIR^E128A^, *At*TIR or *At*TIR^E122A^ at 2 dpi. Bar plots show mean peak areas from three independent biological replicates (each replicate has a different color), with error bars indicating the standard deviation. Constructs labelled with different letters differ significantly (Tukey-HSD, α = 0.05, n = 3).

In addition to being an NADase, TIR-only protein *At*RBA1 can enzymatically degrade double stranded nucleic acid (preferentially RNA) substrates, using different self-association properties and a cysteine catalytic residue (Yu et al., 2022). This produces 2’,3’-cAMP/cGMP stress signals through TIR cyclic nucleotide synthetase activity requiring a triple lysine (KKK) motif for nucleic acid binding (Yu et al., 2022). Members of the conserved TIR-only family possess the conserved catalytic cysteine (C) but lack the triple lysine (KKK) motif, suggesting they do not target DNA or RNA (Figure 1C, Supplementary Figure S1C). A flexible BB-loop motif lying close to the catalytic site of TIR domains supports NAD^+^/ATP substrate binding and *At*RBA1 formation of liquid condensates to increase TIR concentration and NADase specific activity (Li et al., 2023; Song et al., 2024). The predicted BB-loops of *At*TIR and *Hv*TIR differ from *At*RBA1 but share key residues with other conserved TIR-only proteins, including *Nb*TIR, *Os*TIR, *Zm*TIR3 from maize and *Bd*TIR (Figure 1C, Supplementary Figure S1C). To test the function of this predicted intrinsically disordered BB-loop region (Song et al., 2024) in *At*TIR and *Hv*TIR, we replaced residues K75-K80 in *At*TIR and R92-R96 in *Hv*TIR with alanines to produce *At*TIR^bb^ and *Hv*TIR^bb^ mutant variants. The mutants failed to induce cell death although they accumulated to similar levels as the corresponding WT TIR-only proteins in *Nb* transient assays (Figure 1A, B, Supplementary Figure S1A, S1E). Moreover, *At*TIR and *Hv*TIR, but not *At*TIR^bb^ and *Hv*TIR^bb^, formed weak cytoplasmic puncta resembling phase-separated condensates, similar to the positive control *At*RBA1 in non-triggered cells (Supplementary Figure S1D, E (Song et al., 2024)). These results suggest that *At*TIR and *Hv*TIR require an intact NADase catalytic site and BB-loop facilitated assembly to elicit cell death in *Nb*.

Next we analyzed TIR-NADase generated metabolites by transiently expressing *At*TIR, *Hv*TIR or their respective NADase catalytic mutants *At*TIR^E122A^ and *Hv*TIR^E128A^ in *Nb epss* plants and quantifying NAD^+^ hydrolysis products by LC-mass spectrometry (LC-MS). We were not able to directly measure the EDS1 dimer-activating second messengers pRib-AMP/ADP and ADPr-ATP/di-ADPR in plant extracts, as experienced in other studies (Huang et al., 2022; Jia et al., 2022; Bayless et al., 2025). Instead, we measured accumulation of 2’cADPR as a stable biomarker of TIR NADase activity and proposed pRib-AMP/ADP precursor (Wan et al., 2019; Wu et al., 2024; Yu et al., 2024; Bayless et al., 2025). *Hv*TIR and *At*TIR WT proteins produced significantly higher 2’cADPR accumulation compared to the *Hv*TIR^E128A^ and *At*TIR^E122A^ NADase catalytic mutants (Figure 1D). All proteins were expressed, although *Hv*TIR and its catalytic mutant accumulated to lower levels than the *A*tTIR variants (Supplementary Figure S1F). Some plant TIR domains also produce the 3’cADPR isomer in *Nb* transient assays, although typically at lower levels than 2’cADPR (Bayless et al., 2023). We detected 3’cADPR with *Hv*TIR and *At*TIR and a lower 3’cADPR signal with *Hv*TIR^E128A^ and *At*TIR^E122A^ (Figure 1D). These results show that members of the conserved TIR-only family from monocots and dicots produce 2’cADPR and 3’cADPR in *Nb*.

### *Hv*TIR NADase elicited cell death is *EDS1-PAD4-ADR1* independent in barley

To examine *Hv*TIR NADase activity and outputs in a native cell context, we transiently expressed *Hv*TIR-YFP under a maize ubiquitin promoter (*Zm*UBQ) in protoplasts of barley GP (Saur et al., 2019). As a vector-matched negative control, we used a *Zm*UBQ-driven YFP-YFP construct (hereafter YFP). *Hv*TIR, but not the NADase defective mutant *Hv*TIR^E128A^ or YFP induced cell death at 16 h post transfection as measured by reduced luciferase activity reporting a loss of cell viability (Saur et al., 2019), Figure 2A). Protein accumulation was confirmed by immunoblotting (Supplementary Figure S2A). To test the dependency of *Hv*TIR induced cell death on the *HvEDS1-PAD4-ADR1* node, we generated two independent CRISPR/Cas9 homozygous *eds1 pad4* double mutant (*ep-d1* and *ep-*d2) and two independent *adr1* single mutant (*adr1-1* and *adr1-2*) lines in barley GP (Supplementary Figure S2B). All selected barley mutants exhibited normal development and growth compared to GP WT (Supplementary Figure S2B). In luciferase assays of transfected protoplasts, *Hv*TIR, but not *Hv*TIR^E128A^, induced cell death in WT and the *ep-d1, ep-d2, adr1-1* and *adr1-2* mutants (Figure 2A, Supplementary Figure S2C). Therefore, *Hv*TIR-mediated cell death in barley transient over-expression assays also depends on its NADase activity but not EDS1-PAD4-ADR1 signaling.

**Figure 2:**
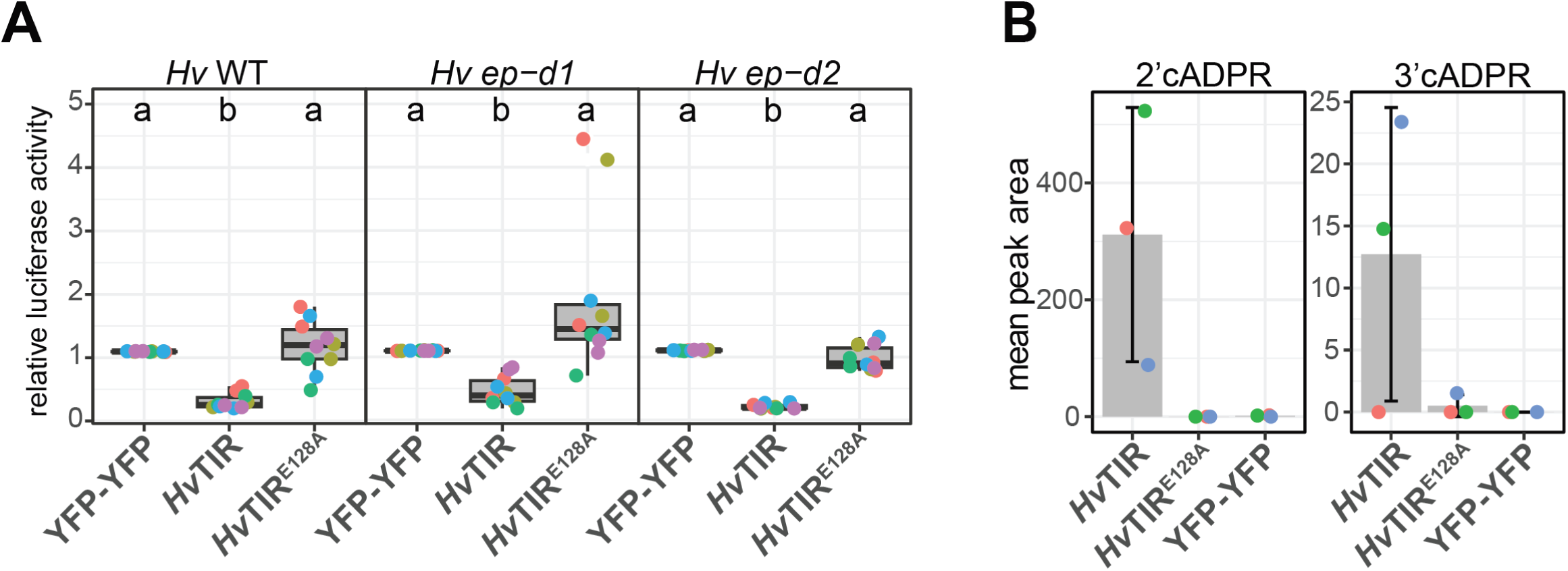
*Hv*TIR induces *EDS1-PAD4*-independent cell death and produces 2’cADPR in barley protoplasts. (A) Relative luciferase activity (RLU) was measured as a proxy for cell death in 7-9 d old barley mesophyll-derived protoplasts of Golden Promise (*Hv* WT) and *eds1-pad4* mutants (*ep-d1*, *ep-d2*). Protoplasts were transfected with YFP -tagged *Hv*TIR, *Hv*TIR^E128A^ or YFP-YFP. Luciferase activity measured ∼16 h post-transfection indicated cell viability and was normalized to luminescence of the YFP -YFP negative control. The experiment was repeated five times independently (n:2 per experiment). Constructs with different le tters indicate significant differences (Kruskal-Wallis test with Nemenyi post -hoc, α = 0.05, n = 10). For boxplots, the center line indicates the median, edges show the 25th and 75th percentiles, and whiskers indicate 1.5 × IQR. (**B**) Quantification via LC-MS of 2’cADPR and 3’cADPR levels in *Hv* WT protoplasts 16 h post-transfection with the same *Hv*TIR, *Hv*TIR^E128A^ and YFP constructs as in (A). Bar plots illustrate the mean peak area obtained from chromatograms from three independent replicates shown by different colors, with error bars representing standard deviation.

To investigate whether *Hv*TIR produces 2’cADPR in barley, GP WT protoplasts were transfected with *Hv*TIR, *Hv*TIR^E128A^ or YFP as a negative control. Protein expression of all constructs was confirmed after 16 h (Supplementary Figure S2D) and samples were analyzed by LC-MS (Figure 2B). *Hv*TIR but not *Hv*TIR^E128A^ and YFP produced 2’cADPR, confirming *Hv*TIR NADase activity in barley. Low levels of 3’cADPR were also detected in all samples with *Hv*TIR but not *Hv*TIR^E128A^ or YFP (Figure 2B). Together, these assays demonstrate that *Hv*TIR is an active NADase in barley protoplasts, producing 2’cADPR together with low levels of 3’cADPR in a catalytic Glutamate dependent manner.

### The Arabidopsis conserved TIR-only clade is necessary for basal immunity to *Hpa*

To assess the contribution of conserved TIR-only proteins to pathogen restriction in *Arabidopsis* Col-0, we generated two homozygous *tir-only* double mutants (*tir-only-d1* and *tir-only-d2*) targeting *AtTIR* and the immune non-responsive TIR-only paralog *AT1G61105* using CRISPR/Cas9 mutagenesis (Supplementary Figure S3A). The *tir-only* mutants developed normally compared to WT plants (Supplementary Figure S3A). Whole-genome sequencing of both *Arabidopsis* mutant lines confirmed presence of the targeted mutations in *AtTIR* and *AT1G61105* (Supplementary Table S1). No other TIR-domain gene was altered except *AT4G23440* (*At*TNP-IIb, TN17 expressing a predicted 942 amino acid protein (Supplementary Figure S3C, Supplementary Data S1, Supplementary Table S1, (Johanndrees et al., 2022)) in both *tir-only* mutant lines. This second-site mutation resulted in an 80 amino acid truncation at the C-terminus of *At*TNP-IIb tetratricopeptide repeat (TPR) domain (Supplementary Figure S3B, Supplementary Data S1). A second *At*TNP gene, *AT5G56220* (*At*TNP-I, TN21), was unaffected (Supplementary Table S1). Previous analyses of CRISPR-targeted TNP mutations in *Nb* and rice revealed no immune related defects (Johanndrees et al., 2022; Wu et al., 2024). We therefore took on the *Arabidopsis tir-only-d1* and *tir-only-d2* lines to test for conserved *TIR-only* mutant phenotypes.

We assessed Arabidopsis *tir-only-d1* and *tir-only-d2* basal immunity phenotypes to the virulent bacterial pathogen *Pseudomonas syringae* pv. *tomato* (*Pst)* strain DC3000 and ETI phenotypes to *Pst* DC3000 expressing TNL RRS1/RPS4-recognized effector AvrRps4 (Birker et al., 2009; Narusaka et al., 2009) or CNL RPS2-recognized effector AvrRpt2 (Bent et al., 1994) (Supplementary Figure S3C). The *tir-only* mutants did not alter bacterial growth compared to WT Col-0 for any *Pst* DC3000 strain. Therefore, the conserved TIR-only genes are dispensable for basal immunity and ETI responses to *Pst* bacteria (Supplementary Figure S3C).

We next examined basal resistance of the *tir-only-d1* and *tir-only-d2* mutants and Col-0 WT to inoculation with a virulent oomycete pathogen *Hyaloperonospora arabidopsidis* (*Hpa*) strain Noco2. Numbers of *Hpa* conidiospores produced on leaves were quantified at 5 dpi (Stuttmann et al., 2011). *Hpa* Noco2 effector ATR1 is recognized by TNL RPP1-WsB in Arabidopsis accession Ws-2, leading to ETI restriction of pathogen colonization (Botella et al., 1998) (Figure 3A). The conserved *tir-only* mutants displayed similar enhanced susceptibility to *Hpa* Noco2 as the Col-0 *eds1-12* mutant (Ordon et al., 2017), indicating that these TIR-only proteins are required for Arabidopsis basal immunity to *Hpa* (Figure 3A). We concluded that this small conserved TIR-only group has a role in Arabidopsis basal immunity to *Hpa* infection which, surprisingly, is not compensated for by other TIR-containing genes.

**Figure 3:**
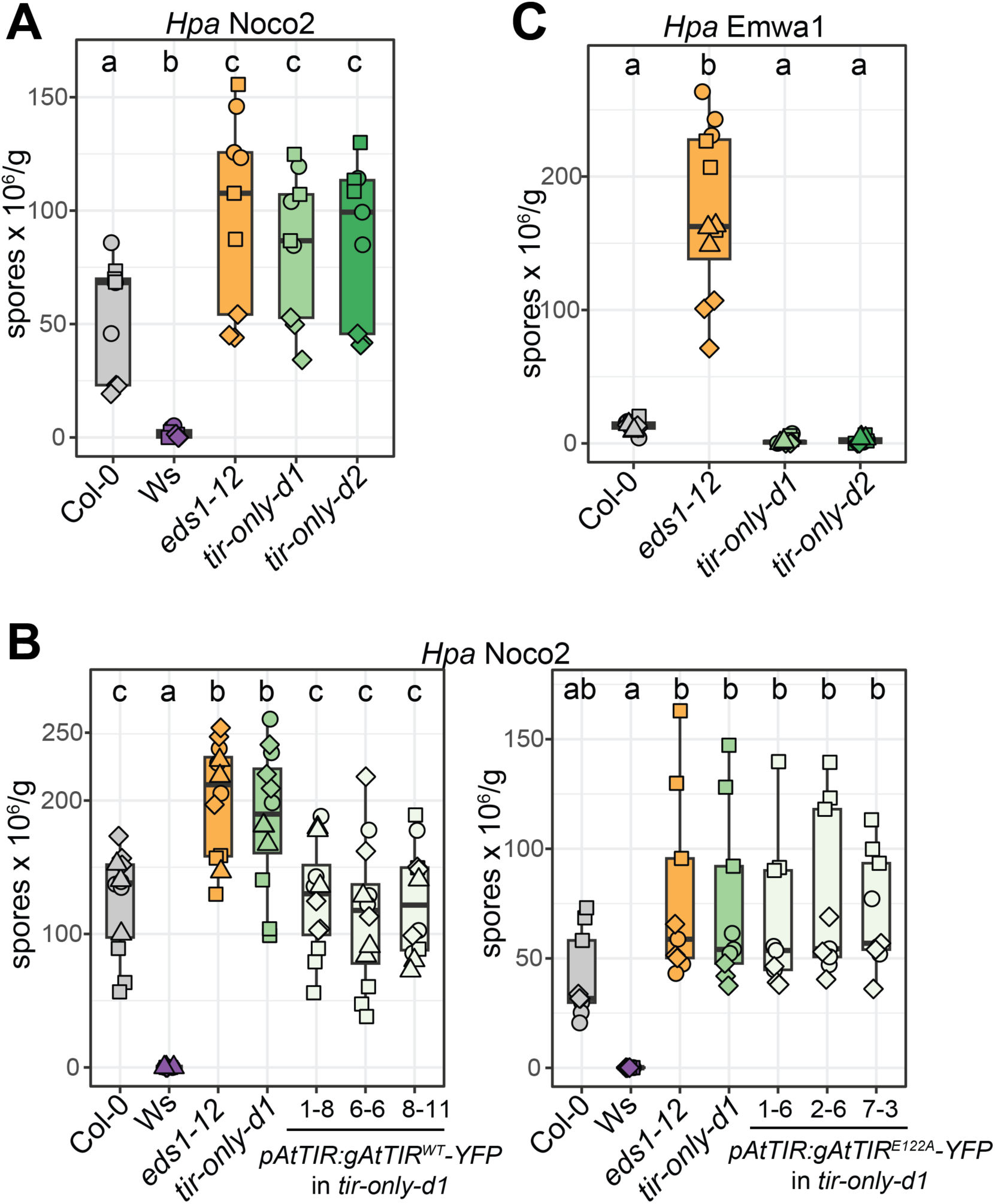
Arabidopsis conserved TIR -only proteins mediate basal immunity to *Hpa*. *Hpa* isolate Noco2 infection phenotypes of 3-week-old plants of ( **A**) Col-0, Wassilewskija (Ws -2), *eds1-12*, *tir-only-d1*, *tir-only-d2*, and (**B**) Independent *At*TIR^WT^ and *At*TIR^E122A^ transgenic lines of *tir-only-d1*, as indicated. Pathogen spores on leaves were counted at 5 or 6 d after spray-inoculation with 4 × 10 ^4^ spores ml−1, normalized to genotype fresh weight (spores x 10 ^6^/g). The experiment was repeated independently three times, each with three technical replicates (n:3, with n=pool of 4 -6 plants). Datapoints with the same shape are from one experiment. Different letters indicate statistically significant difference s (Tukey-HSD, α = 0.05, n = 9-12 or Kruskal-Wallis test with Nemenyi post-hoc, α = 0.05, n = 9). (**C**) *Hpa* isolate Emwa1 infection phenotypes on 3-week-old plants of Col -0, *eds1-12*, *tir-only-d1* and *tir-only-d2*. Pathogen spores on leaves were counted at 5 or 6 d after spray-inoculation as described in (A). The experiment was repeated independently three times, each with three technical replicates (n:3, with n=pool of 4-6 plants). Datapoints with the same shape are from one experiment. Different letters indicate statistically significant differences (Tukey-HSD, α = 0.05, n = 12). For boxplots, the center line indicates the median, edges show the 25th and 75th percentiles, and whiskers indicate 1.5 × IQR.

To test whether NADase activity of the conserved TIR-only proteins is required for *Hpa* Noco2 growth restriction, *At*TIR^WT^ or its catalytic mutant *At*TIR^E122A^, each fused to a C-terminal YFP and driven by the *AtTIR* native promoter, were stably transformed into mutant *tir-only-d1*. Three independent transgenic lines for each construct were selected based on induction of transgene expression following flg22 treatment, a peptide epitope of bacterial flagellin recognized by the LRR receptor kinase FLAGELLIN SENSING 2 (FLS2) (Zipfel et al., 2004) , which was used to assess inducible expression from the native *AtTIR* promoter (Supplementary Figure S3D). While accumulation of *At*TIR^WT^ proteins was readily detected in these lines, *At*TIR^E122A^ proteins were not detected despite their transcriptional induction (Supplementary Figure S3E), suggesting reduced stability of the catalytically inactive TIR-only variant in Arabidopsis. Complementation with *AtT*IR^WT^, but not *At*TIR^E122A^, restored basal resistance to *Hpa* Noco2 (Figure 3B). These results further support a non-redundant role of conserved TIR-only NADase activity in restricting *Hpa* infection.

To determine whether the conserved Arabidopsis TIR-only proteins contribute to TNL mediated ETI to *Hpa*, we inoculated the *tir-only-d1* and *tir-only-d2* mutants with *Hpa* isolate Emwa1 which is recognized by TNL RPP4 in Col-0 WT (van der Biezen et al., 2002). The *tir-only* mutants displayed a similar ETI response to Col-0, whereas *eds1-12* was susceptible (Figure 3C). Taken together, the results show that conserved TIR-only proteins in Arabidopsis are necessary for inducing basal immunity against *Hpa* but are dispensable for basal immunity to *Pst* bacteria and TNL mediated ETI.

### A single barley conserved *TIR-only* gene promotes basal immunity to powdery mildew

The role of *Hv*TIR or functions of *EDS1* and *PAD4* in barley immunity remain unknown. To address this, we generated CRISPR/Cas9-targeted *HvTIR* mutations in the barley cultivar GP, producing two mutant lines (*tir-only-1* and *tir-only-2*, Supplementary Figure S4). Both mutants developed similarly to WT plants (Supplementary Figure S4). To test whether these genes contribute to basal immunity, we infected *tir-only-1* and *tir-only-2* alongside the *eds1-pad4* double mutant lines *ep-d1* and *ep-d2* with virulent strain K1 of the fungal powdery mildew pathogen *Blumeria graminis* f. sp. *hordei* (*Bgh*) (Lu et al., 2016). All mutants displayed increased susceptibility to *Bgh* K1 compared to WT (Figure 4A). In these assays, the resistant cultivar Pallas 01 (P01) which carries a *Bgh* K1 effector-recognizing coiled-coil NLR (CNL) *Mla1* (Kølster et al., 1986) fully restricted pathogen growth (Figure 4A). These results support a co-function of *Hv*TIR with *Hv*EDS1-*Hv*PAD4 in promoting basal immunity to powdery mildew disease.

**Figure 4:**
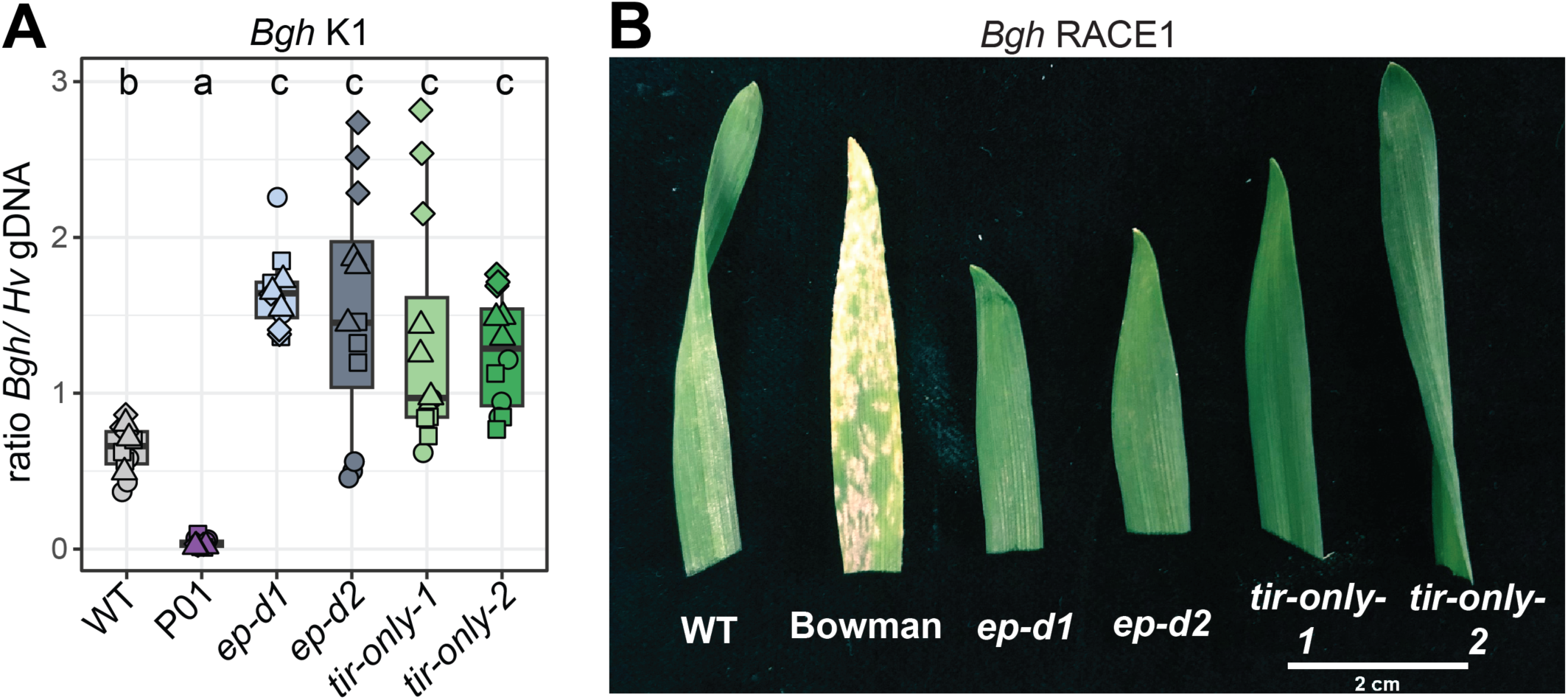
*Hv*TIR is essential for basal immunity to powdery mildew. (**A**) RT-qPCR analysis of *Blumeria graminis* f. sp. *hordei* (*Bgh*) K1 infection in 1.5-week-old barley WT (GP), cultivar Pioneer 01 (P01), *ep-d1*, *ep-d2*, *tir-only-1*, and *tir-only-2* at 5 dpi. Expression levels of *Bgh* K1 *GAPDH* and barley *ACTIN* were used to estimate genomic DNA ratios and thus pathogen load. The experiment was repeated four times independently (indicated by shape) with three technical replicates. Genotypes with different letters above boxplots indicate significant differences (Tukey-HSD, α = 0.05, n = 12). For boxplots, the center line indicates the median, edges show the 25th and 75th percentiles, and whiskers indicate 1.5 × IQR. (**B**) *Bgh* RACE1 infection of barley WT (GP), variety Bowman, *ep-d1*, *ep-d2*, *tir-only-1*, and *tir-only-2* at 7 dpi. For each genotype, 8-9 plants were grown per pot, and one representative leaf from these plants is shown. The experiment was repeated four times independently with consistent results.

We assessed whether *Hv*TIR and/or *Hv*EDS1/*Hv*PAD4 contribute to CNL mediated ETI against *Bgh* isolate RACE1 which is recognized by a CNL *Mla8* allele in GP (Yaeno et al., 2021). Following *Bgh* RACE1 infection, the two *tir-only* and *eds1-pad4* mutant lines were as resistant as WT, while a RACE1 non-recognizing barley cultivar Bowman was susceptible (Leng et al., 2020) (Figure 4B). Therefore, neither *Hv*TIR nor the *Hv*EDS1-*Hv*PAD4 module is required for the CNL MLA8 mediated ETI response in barley.

Put together, our results show that the small group of conserved TIR-only proteins share an essential role in basal resistance against virulent biotrophic filamentous pathogens in Arabidopsis (*Hpa*; Figure 3A) and barley (*Bgh*; Figure 4A). Moreover, the immune contributions of these components seem to be independent of canonical NLR mediated ETI pathways, underscoring their basal immunity function in dicots and monocots.

### Conserved TIR-only proteins promote PTI in Arabidopsis and barley

A PAMP pretreatment enhances resistance to subsequent pathogen infection through activation of PTI (Zipfel et al., 2004; Bohm et al., 2014). In Arabidopsis this defense response is associated with the induction of TIR-domain genes and involves EDS1-PAD4-ADR1 signaling (Pruitt et al., 2021; Tian et al., 2021). Using the *tir-only* mutant lines generated in Arabidopsis and barley, we tested whether conserved TIR-only proteins contribute to immunity against virulent *Pst* DC3000 induced by prior PAMP application. For this we applied the flg22 epitope of bacterial flagellin which is perceived by LRR-receptor kinase (RK) FLAGELLIN-SENSING 2 (FLS2) (Zipfel et al., 2004) or the oomycete nlp20 epitope recognized by receptor-like protein RLP23 (Albert et al., 2015), allowing us to measure conserved TIR-only functions in both RK- and RLP-mediated PTI. Pre-treatment with nlp20 or flg22 enhanced resistance to *Pst* DC3000 in Arabidopsis Col-0 WT but not in the respective recognition-deficient mutants *rlp23-1* and *fls2-17* (Figure 5A). Arabidopsis *tir-only-d1* and *tir-only-d2* mutants were defective in nlp20-induced resistance and partially impaired in flg22-induced resistance, suggesting that the conserved TIR-only proteins stimulate PAMP elicitor-induced protection against *Pst* DC3000 (Figure 5A). Analysis of previous Arabidopsis RNA-seq data revealed that conserved TIR-only *AtTIR* expression was transiently upregulated at 2 h after *Pst* DC3000 inoculation, independently of HopAM1 effector delivery (Figure 5B, (Iakovidis et al., 2016)). These data suggest that Arabidopsis conserved TIR-only proteins have an important role in PTI.

**Figure 5:**
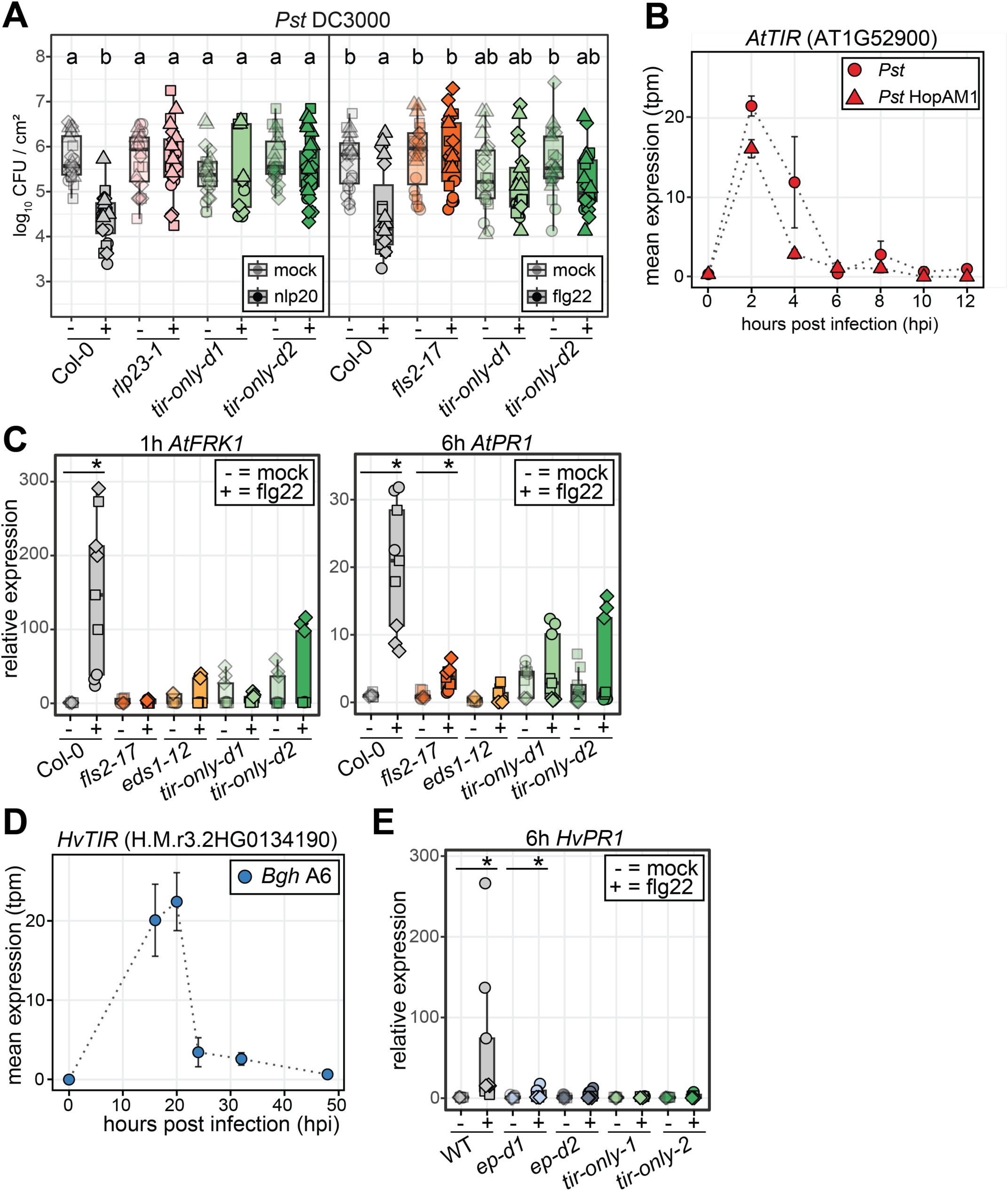
*At*TIR and *Hv*TIR promote defense marker gene induction. (**A**) PAMP elicitor -induced immunity against *Pst* DC3000 was performed in 4/5 -week-old Arabidopsis plants of Col -0, *rlp23-1*, *fls2-17*, *tir-only-d1*, and *tir-only-d2*. Leaves were syringe infiltrated with mock (DMSO), nlp20 (0.5 µM, left panel) or flg22 (0.5 µM, right panel) and challenged with *Pst* DC3000 after 24 h (OD _600_=0.0005). Boxplots show bacterial colonization as log _10_ colony-forming units (CFU) per cm ^2^ at 3 dpi (n:12 biological replicates in four independent experiments comprising 2 leaf discs). Different letters indicate statistically significant difference (Tukey-HSD, α = 0.05, n = 24). Colors indicate genotypes, opacity indicates treatment. Datapoints with the same shape are from one experiment. (**B**, **D**) Induction of Arabidopsis *TIR-only* (*AtTIR*, AT1G52900, panel B) and *Hordeum vulgare TIR -only* (*HvTIR*, H.M.r3.2HG0134190, panel D) genes during pathogen infection. *AtTIR* expression was analyzed using RNAseq data (SRP075162) from Arabidopsis Col -0 infected with *Pst* strain D28E or *Pst* transformed with HopAM1 at 0 -12 hpi. *HvTIR* expression was analyzed using RNAseq data (SRP111697) from barley GP infected with *Bgh* isolate A6 at 0 -50 hpi. Expression values are normalized to transcripts per million (tpm) with error bars representing the standard error of the mean. ( **C, E**) Induction of *FRK1* and *PR1* defense genes in Arabidopsis ( *AtFRK1*, *AtPR1*, panel C) and *PR1* in barley (*HvPR1*, panel E) after flg22 treatment (0.5 µM for Arabidopsis, 1 µM for barley). RT-qPCR was conducted on 4-week-old Arabidopsis (Col-0, *fls2-17*, *eds1-12, tir-only-d1*, *tir-only-d2*) and 2-week-old barley (WT, *ep-d1*, *ep-d2*, *tir-only-1*, *tir-only-2*) plants at 1 and 6 h post treatment. Relative expression (RQ) was calculated against mock treatments for each time point, normalized to the housekeeper genes *UBI5* in Arabidopsis and *UBIQUITIN* in barley. Boxplots show RQ values with different shapes indicating independent experiments. Asterisks indicate significant differences between mock and flg22 treatments (Wilcoxon signed -rank test, p < 0.05, n = 9). For boxplots, the center line indicates the median, edges show

Typical hallmarks of PTI include production of extracellular reactive oxygen species (ROS), phosphorylation of mitogen-activated protein kinases (MAPKs), cytosolic Ca²⁺ influx and transcriptional reprogramming (Boudsocq et al., 2010; Xu et al., 2022). Flg22-induced ROS production and MAPK phosphorylation were unaffected in the Arabidopsis *tir-only-d1* and *tir-only-d2* mutants (Supplementary Figure S5 A-C). However, Flg22-induced expression of the early defense marker gene *FLG22-INDUCED RECEPTOR-LIKE KINASE 1 (FRK1)* (1 h after flg22 treatment) and late defense marker *PATHOGENESIS-RELATED 1* (*PR1*) (6 h after flg22 treatment) was markedly reduced in *tir-only-d1* and *tir-only-d2* compared with Col-0 WT plants (Figure 5C).

In barley, we performed time-resolved analysis of *HvTIR* expression using publicly available RNA-seq dataset (Hunt et al., 2019). This revealed *HvTIR* upregulation at 16–20 h after infection with the virulent fungal pathogen *Bgh* isolate A6 (Figure 5D,(Hunt et al., 2019)), corresponding to an early stage of haustorium formation in epidermal cells (Yamaoka et al., 2000). Similar to Arabidopsis, flg22 treatment of barley triggers a ROS burst, MAPK phosphorylation, and transcriptional reprogramming (Proels et al., 2010; Boyd et al., 2013). We found that flg22-induced ROS production and MAPK phosphorylation were unaffected in the barley *tir-only-1, tir-only-2* and *ep-d1*, *ep-d2* mutants (Supplementary Figure S5 D–F). However, all mutants failed to induce expression of the *HvPR1* defense marker at 6 h after flg22 treatment when compared to WT (Figure 5E). These data show that conserved TIR-only proteins are required for induced defense gene expression following flg22 perception in Arabidopsis and barley, thus highlighting a potentially fundamental TIR-only role in PTI associated transcriptional responses in diverse seed plants.

### Flg22 induced *HvTIR* and *HvEDS1-PAD4* dependent regulation of defense pathways in barley

As transcriptional responses to PTI activation remain poorly characterized in barley we performed RNA-seq analysis of flg22-triggered gene expression changes. In a pilot study, we measured the expression of *HvEDS1*, *HvPAD4*, and *HvADR1* in barley GP at 30 min, 1 h and 6 h after mock or flg22 treatment (Supplementary Figure S6A). Barley *EDS1* and *PAD4* were upregulated by flg22 at 1 h (Supplementary Figure S6A). Curiously, *HvADR1* expression decreased at 30 min and 6 h but remained unchanged at 1 h (Supplementary Figure S6A). The 1h time point was therefore selected for transcriptome analysis.

RNA-seq was performed on leaf disc samples from 1h flg22- vs mock-treated barley WT, *ep-d1*, *ep-d2*, *tir-only-1*, and *tir-only-2* plants. Principal component analysis (PCA) revealed a clear separation between mock- and flg22-treated samples, explaining 39% of the variance (Figure 6A). Differential expression analysis (log_2_FC > 1) identified genes whose expression changed specifically in response to flg22 compared to mock in each genotype. A Venn diagram highlights differentially expressed genes (DEG; 75 up- and 20 down-regulated) in WT versus all mutants, indicating transcriptional changes robustly associated with *Hv*TIR and *Hv*EDS1*-Hv*PAD4 signaling (Figure 6B, Supplementary Figure S6B). Gene Ontology (GO) analysis (hypergeometric testing, FDR < 0.05) of the 75 flg22-responsive genes that were upregulated in WT but not in the *eds1-pad4* and *tir-only* mutants (Figure 6B, Supplementary Table S2, Supplementary Figure S7) showed enrichment for kinase activity, ATP binding, calcium signaling, and transcription (Supplementary Figure S8A, Supplementary Table S3). Gene annotations (Mascher et al., 2021) further identified potential kinases and transcription factors (Figure 6C). A member of the calcium-dependent phospholipid-binding copine family was among genes upregulated in WT but not the mutants (Figure 6C). This is notable because copine (BONZAI) orthologs have been characterized as Ca^2+^ and pathogen-regulated defense suppressors in Arabidopsis, wheat and rice (Yang and Hua, 2004; Lee and Mcnellis, 2009; Yin et al., 2018; Zou et al., 2018). No commonly downregulated genes were found specifically in the mutants, whereas 20 genes specifically downregulated in WT were enriched for proteolysis based on GO annotations (Supplementary Figure S6C).

**Figure 6:**
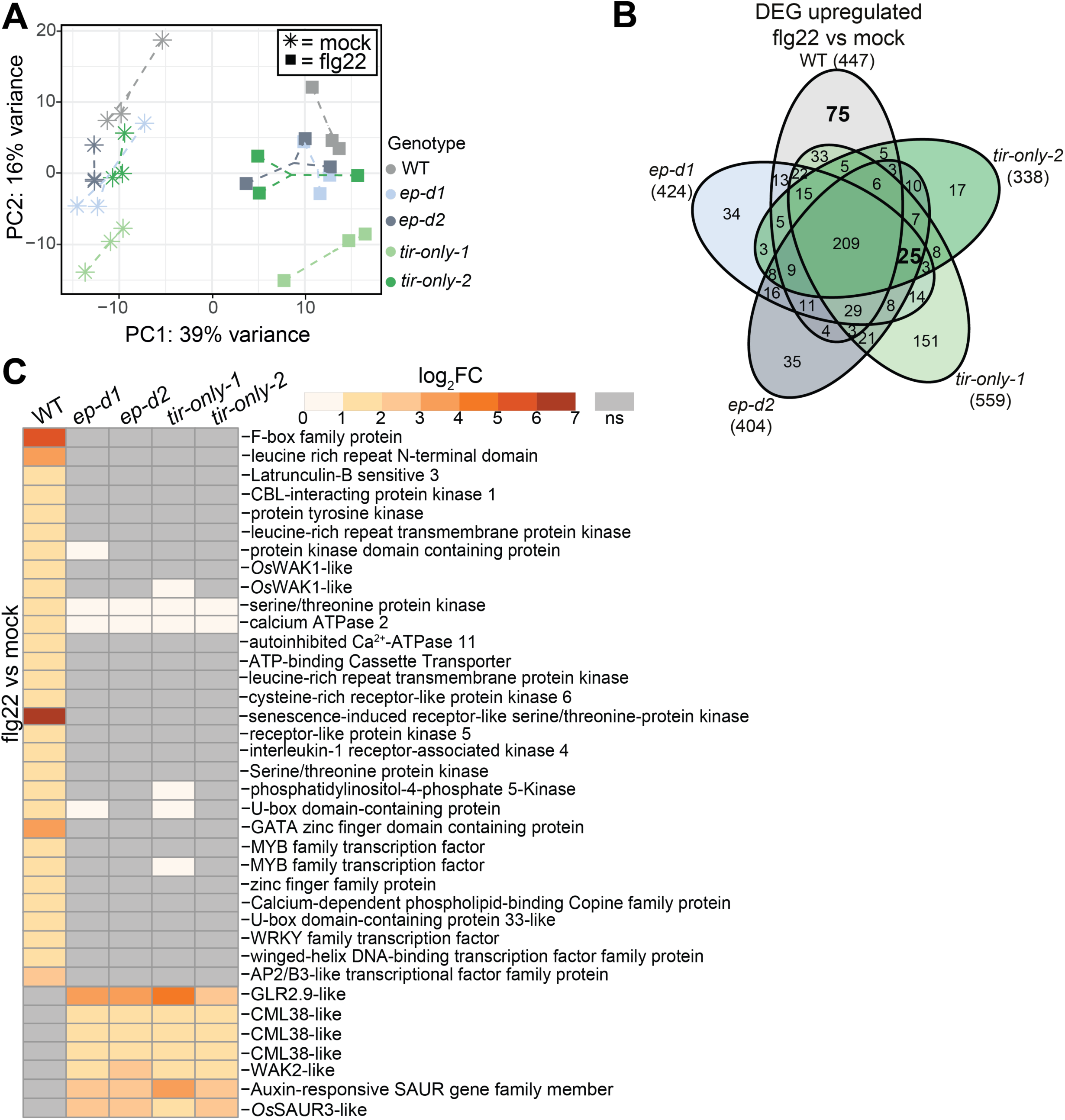
*HvTIR*, *EDS1* and *PAD4* regulate early transcriptional changes following flg22 treatment in barley. (**A**) Principal Component Analysis (PCA) of the top 500 most variable genes based on expression levels in 2-week-old barley WT (GP), *ep-d1*, *ep-d2*, *tir-only-1*, and *tir-only-2*, 1 h after mock or 1 µM flg22 treatment. Shapes denote treatments and colors represent genotypes. The plot displays the first two principal components. (**B**) Differential gene expression analysis comparing flg22 and mock treatments in barley WT, *ep-d1*, *ep-d2*, *tir-only-1*, and *tir-only-2*. Genes with an adjusted p -value < 0.05 and log_2_ fold-change > 1 were deemed significant. 75 DEG exclusive to WT and 25 DEG shared by all mutants are marked in bold. (**C**) The heatmap displays log_2_ fold-change expression of genes identified in (**B**) associated with significant GO terms - ATP binding, kinase activity, transcriptional responses, auxin responses and calcium ion binding across the genotypes. Non -significant (ns) changes are shown in grey.

We identified a set of 25 genes as upregulated in the *eds1-pad4* and *tir-only* mutants but not in WT after flg22 treatment (Figure 6B, Supplementary Table S4, Supplementary Figure S9A), implying these genes are suppressed by TIR-stimulated EDS1-PAD4 signaling in barley PTI. GO term enrichment for the genes (hypergeometric testing, FDR < 0.05) revealed significant representation of auxin responses and calcium regulation (Supplementary Figure S9B, Supplementary Table S5). In this expression group, three genes annotated as *WALL ASSOCIATED RECEPTOR KINASE 2-LIKE (WAK2)* and *CALMODULIN-LIKE 38* (*CML38*) are associated with calcium ion binding (Figure 6C, (Mascher et al., 2021)). Additional calcium-related genes, including a second *CML-38* like and a *GLUTAMATE RECEPTOR 2.9-LIKE* (*GLR 2.9-like*) gene (Bjornson et al., 2021; Wang et al., 2025) were also upregulated in the mutants but not WT, although they were not identified in the GO enrichment analysis (Figure 6C, Supplementary Figure S9B). The barley RNA-seq data show that *HvTIR* and *EDS1-PAD4* promote flg22-induced transcriptional defenses at 1 h while simultaneously repressing a subset of calcium- and auxin-related genes. This suggests an important role of the barley TIR/EDS1-PAD4 node in balancing immune signaling with growth.

## Discussion

Here we examined the genetic, biochemical and immune signaling functions of a small conserved group of TIR-only proteins in Arabidopsis and barley, as representative dicot and monocot species. We established that *At*TIR and *Hv*TIR are pathogen-responsive NADase enzymes (Figure 3) with a capacity to produce ribosylated cyclic nucleotides (Figure 1D, 2B) and elicit cell death when transiently over-expressed in *Nb* (Figure 1A,B) and, in the case of *Hv*TIR, barley protoplasts (Figure 2A). Although we could not detect pRib-AMP/ADP molecules directly *in vivo*, the mutational (Figure 1A-C) and metabolite profiling data (Figure 1D, 2B) are consistent with members of this small conserved TIR-only protein group generating pRib-AMP/ADP signals to stimulate transcriptional defense through the EPA node. Strikingly, Arabidopsis and barley conserved *tir-only* mutants displayed similar basal immunity defects to filamentous virulent pathogen infections, respectively downy mildew (*Hpa*) and powdery mildew (*Bgh*), but were dispensable for ETI (Figure 3 and 4). Also, flg22 PAMP elicitor pre-treatments of conserved *tir*-only mutants in both species did not affect early ROS and MAPK signaling pathways (Supplementary Figure S5) but did compromise PAMP-triggered defense gene induction (Figure 5C,E), as well as flg22-induced immunity to *Pst* DC3000 bacterial infection in Arabidopsis (Figure 5A). Our comparative analysis highlights a fundamentally important role of conserved TIR-only proteins in driving basal immunity in dicots and monocots.

The equivalent phenotypes of barley CRISPR *tir-only-1* and *tir-only-2* compared to the *ep-d1* and *ep-d2* mutants (Figure 4, 5E) underscore *Hv*TIR cofunction specifically with the EPA module, as observed in rice (Wu et al., 2024). Similar penetrance of Arabidopsis *tir-only-d1* and *tir-only-d2* compared to *Ateds1-12* (which abolishes all TIR immune signaling, (Lapin et al., 2020)) (Figure 3, 5C) indicates that loss of these two conserved TIR-only members is not compensated for by the ∼ 160 other TIR-domain genes (Yue et al., 2012; Bayless et al., 2025). We think we can firmly rule out that a second-site mutation in *At*TNP-IIb (*AT4G23440*, TN17) (Supplementary Figure S3B) is causal for failure of *tir-only-d1* and *tir-only-d2* plants to mount PTI defense. TNPs likely have an ancient origin (Gao et al., 2022; Johanndrees et al., 2022; Liu et al., 2023) but did not contribute measurably to immunity in *Nb* and rice (Johanndrees et al., 2022; Wu et al., 2024). Also, *At*TNP-derived TIR domains failed to elicit cell death or 2’cADPR production in *Nb* transient assays (Bayless et al., 2025). Moreover, our analysis of *tir-only-d1* stable transformants expressing native promoter-controlled *At*TIR or its catalytic mutant *At*TIR^E122A^, showed full *At*TIR NADase-dependent rescue of the *tir-only-d1 Hpa* basal immunity defect (Figure 3B). Hence, conserved TIR-only proteins in Arabidopsis, and potentially other dicots, appear to have a unique function in PTI associated defense gene induction. Their key role in PTI signaling could explain retention of the TIR-only group in monocot lineages which have otherwise dispensed with TIR-domain immunity genes, apart from the TNPs.

We considered which characteristics of the conserved TIR-only group might distinguish them from other TIR-only (such as RBA1), TIR-NB and TNL genes present in Arabidopsis and dicots in general (Yue et al., 2012). The conserved TIR-only enzymes could be especially effective in potentiating and spreading immune defense signals, as observed with a biotrophic filamentous pathogen such as *Hpa* tested here (Figure 3). This notion is supported by known contributions of the EPA node in spatial defense reinforcement around Arabidopsis CNL mediated ETI sites (Lapin et al., 2020) and role of conserved TIR-only stimulated EPA pathways in Arabidopsis and *Nb* stomatal immunity (Wang et al., 2024; Wang et al., 2024). The conserved TIR-only enzymes might be particularly ‘quick off the mark’ in response to PAMP and DAMP stimulation due to a low-level capacity already in unstimulated cells to form NADase-competent puncta (Supplementary Figure S1D) coupled with their pathogen-responsive transcriptional induction (Figure 5B,D (Iakovidis et al., 2016; Hunt et al., 2019).

Emerging evidence that plant TIR-domain proteins can act at different stages and cellular locations in defense (Jacob et al., 2023; Tang et al., 2023) aligns with a non-redundant conserved TIR-only contribution in Arabidopsis immunity. Arabidopsis TNL Suppressor of ADR1-L2 1 (SADR1) stimulates late-stage RLP23-triggered PTI responses, including *PR1* expression and restriction of bacterial pathogen growth (Jacob et al., 2023), whereas SM-binding competent EDS1-PAD4 and ADR1 mediate early- and late-phase PTI (Fliegmann et al., 2025). The Arabidopsis *tir-only-d1* and *tir-only-d2* mutants were defective in late-stage PTI outputs (Figure 5C) and thus might enable SADR1 and other TIR-domain enzymes to boost defense beyond infection sites. Notably, Tang et al. (2023) detected expression of multiple TIR-domain genes in vascular bundle sheath cells distal from sites of fungal colonization, suggesting a relay of TIR enzymes and potential products assist systemic defense propagation in leaves.

It is plausible that the conserved TIR-only enzymes have a particular in vivo mode of activation, substrate preference and/or ribosylated nucleotide profile. A recent *At*TIRome study identified TIR enzymes with different 2’cADPR accumulation and cell death properties in *Nb* transient assays, influenced by variation in the TIR BB-loop lying close to the catalytic site (Bayless et al., 2025). Dicot and monocot conserved TIR-only members possess a variant BB-loop (Figure 1C) which could thus alter their biochemical activities. However, in *Nb* assays *At*TIR and *Hv*TIR enzymatic profiles were similar to many other tested *At*TIR-domains (Figure 1D) (Bayless et al., 2025). Our detection of 2’cADPR (as a potential pRib-AMP precursor) with 3’cADPR as a minor product is similar to the profile of *Bd*TIR (Wan et al., 2019; Bayless et al., 2023). In bacteria, TIR-generated 3’cADPR is an important anti-phage immune signal (Leavitt et al., 2022; Manik et al., 2022) and TIR NADase effector product (Eastman et al., 2022) with yet unknown activity in plants (Huang et al., 2022; Manik et al., 2022). In another recent study of maize conserved TIR-only proteins, metabolite profiling of WT and catalytic mutants revealed their capacity to produce 2’cADPR as well as 2’,3’-cNMPs (Zhang et al., 2025). Unlike *At*RBA1 and a TIR isolated from flax rust resistance TNL L7, monocot TIR-only sequences (including those of maize and *Bd*) possess a conserved cysteine for cyclic synthase activity but lack an intact ‘KKK’ site for binding nucleic acid substrates (Supplementary Figure S1C). This raises the possibility that conserved TIR-only proteins generate 2’,3’-cNMP molecules through a distinct substrate preference or catalytic mechanism.

A limitation to TIR transient over-expression in *Nb* is that it is unlikely to reflect the native cell contexts and biochemical micro-environments in which TIR-only proteins operate. Plant TIR enzymes can be modulated by phosphorylation (Li et al., 2025) and Ca^2+-^responsive suppressors such as RESISTANCE OF RICE TO DISEASES1 (ROD1) in rice (Wu et al., 2024) and BONZAI-family copines in Arabidopsis, *Nb*, rice and wheat (Yang and Hua, 2004; Lee and Mcnellis, 2009; Yin et al., 2018; Zou et al., 2018) to balance immunity with growth. We also found that native promoter-controlled *At*TIR protein mutated at its ‘Glu’ catalytic residue failed to accumulate in Arabidopsis stable transgenic lines (Supplementary Figure S3E), whereas its level was similar to WT *At*TIR in *Nb* transient assays (Supplementary Figure S1A, E). While this remains unexplained, it suggests a further level of TIR-only protein regulation in the native sub-cellular context. The diversity of TIR-domain assemblies generally (Li et al., 2023) and TIR catalyzed molecules in bacterial anti-phage defense (Hochhauser and Sorek, 2026), suggest further TIR mechanisms and enzymatic outputs remain to be discovered in plants. Because monocots lack the EDS1-SAG101-NRG1 signaling branch, it remains possible that conserved monocot and dicot TIR-only proteins differ in their broader metabolite outputs. Future developments in LC-MS based ribosylated nucleotide profiling might resolve which molecules predominate in plant TIR-stimulated EPA signaling whose chief role in dicot and monocot immunity is in transcriptional potentiation of defenses, not elicitation of host cell death (this study) (Lapin et al., 2020; Wang et al., 2024; Wu et al., 2024).

Our barley RNA-seq analysis at 1h post flg22 treatment provides further clues to the dynamics of TIR-stimulated EPA-dependent transcriptional defense in PTI, independently of ETI. By categorizing up- and down- DEG in WT that were absent in all *tir-only* and *eds1 pad4* mutant lines we could extract gene expression changes robustly associated with the TIR-EPA module (Figure 6). While the 1h time point is a transcriptional snap shot of barley PTI responses, it emphasizes the importance of Ca^2+^ sensing and Ca^2+^ homeostasis in TIR-EPA controlled defense (Figure 1C). Especially striking was induction of copine family orthologs (Supplementary Table S7) in the barley WT response which was lost in the mutants (Figure 6), suggesting broad significance of these Ca^2+^ and phospholipid binding homeostatic regulators not only in suppressing TIR-based signaling in healthy plants, as found in other studies (Wang et al., 2024; Wu et al., 2024), but also in dampening immune over-activation in PAMP-triggered tissues. Similarly informative is observed deregulation of genes with Ca^2+^ related functions (such as *WAK2*, *CML38*, and *GLR2.9*) specifically in the mutants (Figure 6). Together, these expression trends point to an important contribution of TIR-EPA signaling in barley to a Ca^2+^-responsive, balanced mobilization of PTI defense to maintain growth.

## Materials and methods

### Plant materials and growth conditions

*Arabidopsis thaliana* Col mutants *eds1-12*, *rlp23-1* and *fls2-17* were previously described by (Zipfel et al., 2004; Albert et al., 2015; Ordon et al., 2017). Arabidopsis plants were grown in a growth chamber under conditions of 10h light and 14h dark cycles, with a light intensity of approximately 150 μmol/m²/s, at 22 °C and 65 % humidity. Assays were performed on 4-week-old Arabidopsis plants. Barley cultivars were maintained at 19 °C with 70% relative humidity under a 16h light and 8h dark photoperiod. Protoplast assays were performed on 7-day-old barley plants, while immune assays were performed on 2-week-old plants. *Nicotiana benthamiana* was grown in a greenhouse on a 16h light and 8h dark cycle, and 4- to 5-week-old plants were used for immune assays. *N. benthamiana* mutant *eds1 pad4 sag101a sag101b* (*epss*) was previously described in (Lapin et al., 2019).

### Generation of expression vectors

TIR-only coding sequences, excluding stop codons, were amplified from Arabidopsis Col-0 cDNA using TOPO or BP cloning oligonucleotides (Supplemental Table S8). *Hv*TIR has been published previously (Johanndrees et al., 2022). Amplification was performed using Phusion (NEB) or PrimeStar HS (Takara Bio) polymerases, and sequences were cloned into pENTR/D-TOPO or pDONR221 vectors and verified by Sanger sequencing. Site-directed mutagenesis was used to introduce mutations using specific oligonucleotides (Supplemental Table S8). Sequences were recombined into the pXCSG-GW-YFP expression vector using LR Clonase II (Life Technologies), with correct insertion confirmed by restriction enzyme digests. For *At*TIR complementation studies in *tir-only* mutants, approximately 3 kb of the upstream regulatory region of the Arabidopsis *AtTIR* gene was amplified using high-fidelity polymerases, Phusion (NEB) and PrimeStar HS (Takara Bio), with specific oligonucleotides (Supplemental Table S8). The integrity and size of the PCR products were confirmed by agarose gel electrophoresis, and the purified upstream regions were ligated into pXCSG-GW-YFP vector, replacing the 35S promoter.

### Transient protein expression and cell death assays in *Nicotiana benthamiana* and barley

*Agrobacterium tumefaciens* strains GV3101 pMP90RK containing the plasmid of interest and strains expressing the viral DNA silencing repressor p90 and the RNA silencing repressor Cucumber Mosaic Virus 2b (Koizumi et al., 2017) were incubated in induction buffer (10 mM MES pH 5.6, 10 mM MgCl-2, 150 nM acetosyringone) for 1h in the dark at room temperature. Protein samples for immunoblot assays were collected 2 days post-infiltration (dpi), and macroscopic cell death was documented with a camera at 3 dpi. Two days after infiltration, six 8 mm leaf discs from infiltrated *Nb* leaves were washed with ddH_2_O for 30 min at room temperature and then placed in a 24-well plate with ddH_2_O. Conductivity was measured immediately and after 6h of incubation at room temperature using a Horiba Twin Model B-173 conductometer. Barley protoplast cell death was assessed by luciferase activity as a proxy for cell viability following (Saur et al., 2019). cDNA from *HvTIR*, its catalytic mutant, and YFP were expressed under a strong ubiquitin promoter in a pUBI vector (Shen et al., 2003). Protoplasts of 7-9 days old barley cultivar Golden Promise and its mutants at OD_600_ 0.55 were transfected with 5 µg of luciferase reporter plasmid and 10 µg of TIR plasmids. For measurements of small molecule production and protein accumulation, the volumes were increased tenfold. Protoplasts were incubated at 21 °C for 16h. Harvesting included centrifugation at 1000xg and lysis in 200 µl of 2x cell culture lysis reagent (Promega, E1531). Luciferase activity was measured by mixing 50 µl of lysate with 50 µl of luciferase substrate (Promega, E1501) and recording light emission for 1 second per well using a microplate luminometer (Centro, LB960). Relative luciferase activity was normalized to the YFP sample, which was set to 1.

### MAPK assay in Arabidopsis and barley

Leaves from 4-6 week-old Arabidopsis were syringe-infiltrated with water (mock) or 0.5 µM flg22 (Biomatik; synthesized peptide QRLSTGSRINSAKDDAAGLQIA). After 15 min, the leaves were harvested and snap-frozen in liquid nitrogen. The MAPK assay in barley was conducted as previously described (Scheler et al., 2016). Leaf discs were taken from the upper part of the second leaves of 14-day-old barley plants, incubated in 2 ml of water per well for 16h in 24-well plates, then transferred to fresh water for 30 min. Leaf discs were then elicited with 1 µM flg22 or mock (water) for 15 min, collected, and snap-frozen in liquid nitrogen. Activated MAPKs were detected using a phospho-specific p44/42 antibody (Cell Signaling Technology, #9102) at a 1:5000 dilution, as described by (Willmann et al., 2014).

### Immunoblot analysis

To assess protein accumulation, four 1 cm leaf discs of *Nb* were collected per sample at 2 dpi, and barley protoplasts were harvested 16 h after transfection and ground in liquid nitrogen. *Nb* samples were resuspended in 2x Laemmli buffer (4% SDS, 20% glycerol, 0.004% bromphenol blue, 0.125 M Tris-HCl pH 6.8, 10% β-mercaptoethanol) and boiled at 95°C for 10 minutes. Barley protoplasts were resuspended in plant protein extraction buffer (200 mM Tris-HCl pH 7.5, 150 mM NaCl, 10 mM EDTA, 10% glycerol, 12 mM DTT, 1x plant protease inhibitor cocktail, 1 % IGEPAL), centrifuged, diluted with 2x Laemmli buffer, and boiled at 95 °C for 10 min. Protein extracts were resolved on an Any kD precast sodium dodecyl sulfate polyacrylamide gel (bioRad) and transferred to a nitrocellulose membrane using the Trans Blot Transfer Kit (Bio-Rad, 1704270). Tagged proteins were detected with α-GFP antibodies (1:2500, Roche, 11814460001) or α-MYC (1:5000, Sigma, M4439), followed by HRP-conjugated secondary antibodies (Cell Signaling Technology, #7076). The signal was detected using Clarity™ or Clarity Max™ Western ECL Substrate (Bio-Rad, 1705061 and 1705062) and captured using the Bio-Rad ChemiDoc™ XRS+ detection system.

### Liquid Chromatography Tandem-mass Spectrometry (LC-MS/MS) metabolite measurement

For barley assays, Golden Promise protoplast pellets transfected with *Hv*TIR, *Hv*TIR^E128A^ and YFP were stored at -80 °C. Samples were resuspended in 100 µL of 50% methanol with 1% formic acid, heated at 95 °C for 5 min, sonicated for 10 min, and centrifuged at 17,000g for 5 min at 4 °C. Supernatants were subjected to LC-MS analysis on a Nexera XR HPLC (Shimadzu) coupled to an LCMS-8060 triple quadrupole mass spectrometer using a Synergi Fusion-RP 80 Å column (4 µm, 150 × 4.6 mm; Phenomenex). A 10 µL injection was separated at 0.3 mL min⁻¹ over a 20 min gradient: 0 min, 5% B; 0-5 min, 5% B; 5-8 min, 98% B; 8-15 min 98% B; 15-15.1 min, 5% B using 0.1% formic acid and acetonitrile as mobile phases. Metabolites were detected by scheduled multiple-reaction monitoring (MRM) in positive mode using ion transitions optimized with 2′cADPR (C406, BioLog) and 3′cADPR (C404, BioLog) standards. Data acquisition and analysis were performed using LabSolutions LCMS v5.118 and LabSolutions Postrun, with quantification based on peak integration.

For *N. benthamiana* assays, coding sequences of *Hv*TIR, *At*TIR and their catalytic mutants were expressed from 35S-driven C-terminal 4×MYC (Gateway™ pEarly 101, oligos listed in Supplementary Table S8) fusions in *Nb epss* plants via Agrobacterium GV3101 infiltration (OD600 = 0.5). At 36 hpi, four 8 mm leaf discs from three independent plants were collected, flash-frozen, ground in liquid nitrogen, and extracted in 200 µL of 70 % methanol for 2 h at 4 °C. Extracts were centrifuged and stored at –80 °C before UPLC–MS/MS analysis. Compounds were analyzed on a QTRAP 6500 mass spectrometer (AB Sciex) coupled to an Agilent 1290 UHPLC using an XSelect HSS T3 column (100 × 3.0 mm, 2.5 µm; Waters). The mobile phases were 2 mM ammonium acetate (A) and methanol (B); the flow rate was 0.4 mL min⁻¹, and the gradient was: 0–3 min, 1 % B; 3–9 min, 1–80 % B; 9–9.5 min, 95 % B; 9.5–11 min, 95 % B; 11.1–15 min, 1% B. Analytes were detected in MRM mode with identities confirmed by comparison of mass and retention time with 2′cADPR and 3′cADPR standards. Data were acquired with Analyst 1.6.3 and processed using MultiQuant 3.0.2.

### Hyaloperonospora arabidopsidis growth assays in Arabidopsis

To assess *Hpa* Noco2 and *Hpa* Emwa1 growth, six pots of 3-week-old Arabidopsis plants were spray-inoculated with a spore suspension at a concentration of 4 x 10⁴ spores/ml, as previously described in (Stuttmann et al., 2011). Spore counts were performed five to six days after infection using a Neubauer chamber (Carl Roth). Spore density was then calculated and expressed as spores per gram of plant material.

### *Blumeria graminis* f.sp. *hordei* growth assay in barley

To analyze the *Bgh* growth, the protocol described by (Weßling and Panstruga, 2012) was followed with modifications adapted for barley. Three *Bgh* infected barley plants were used to spread spores on four uninfected, one-week-old barley plants. To assess fungal load of *Bgh* K1 (Jones et al., 2016), the first 5 cm of the leaf tip of the inoculated barley was collected immediately after infection and at 5 dpi, then snap frozen in liquid nitrogen. Samples were lysed in CTAB buffer (2 % cetyl trimethylammonium bromide, 100 mM TRIS-HCl, 1.4 M NaCl, 20 mM EDTA) and DNA was extracted using phenol/chloroform/isoamyl alcohol (25:24:1). DNA concentration was adjusted to 50 ng/µl and analyzed by real-time quantitative PCR (RT-qPCR) using iQ SYBR Green Supermix (BioRad) to quantify fungal DNA relative to plant DNA according to (Weßling and Panstruga, 2012). The primers used are listed in Supplementary Table S8. To assess *Bgh* RACE1 (Lu et al., 2016) growth, one week after infection pictures were taken with a camera.

### *Pseudomonas syringae* (*Pst*) infection and priming in Arabidopsis

*Pseudomonas syringae* pv. *tomato* DC3000 (*Pst* DC3000, (Hinsch and Staskawicz, 1996) was syringe-infiltrated into four- to five-week-old Arabidopsis plants at an OD_600_ of 0.0005 in 10 mM MgCl_2_. Bacterial titers were measured at day 0 and three days after infiltration, as previously described by (Lapin et al., 2019). For priming experiments, leaves were syringe-infiltrated with 0.5 µM flg22 or nlp20 (Genscript, synthesized peptide from *Hyaloperonospora parasitica* AIMYSWYFPKDSPVTGLGHR) in water, or water alone as mock control, 24h prior to *Pst* DC3000 infiltration. *Pst* DC3000 was then infiltrated into the same spots on the leaves 24h after the initial treatment.

### Analysis of publicly available immune-related RNA-seq datasets

Raw RNA sequencing data for Arabidopsis thaliana Col-0 infected with *Pst* expressing HopAm1 or empty vector (EV) (SRP075162 (Iakovidis et al., 2016)) and for barley cultivar Golden Promise infected with *Bgh* isolate A6 (SRP111697 (Hunt et al., 2019)) were retrieved from the NCBI Sequence Read Archive (SRA) using the SRA Toolkit (v2.10.0) (SRA Toolkit Development Team, https://github.com/ncbi/sra-tools). An initial quality assessment of the raw sequencing data was performed using FastQC. Trimmomatic (v0.38) was used to remove residual adapter sequences using parameters such as LEADING:5, TRAILING:5, SLIDINGWINDOW:4:15, MAXINFO:50:0.8, and MINLEN:36 (Bolger et al., 2014). Quantification of reads was performed using Salmon (v1.4.0) with specific options for single end data (--fldMean=150 --fldSD=20) and paired end data (--validateMappings --gcBias) (Patro et al., 2017). The reference genomes used were TAIR10 for Arabidopsis and IBSCv2 for barley. The tximport package (v1.22.0) was used to convert read counts to transcripts per million (TPM) (Soneson et al., 2016). The resulting TPM values were then normalized to z-scores.

### RNA extraction and RT-qPCR

To measure transcript levels of genes, 4-6 week old Arabidopsis plants were treated with mock (water) or 0.5 µM flg22 (Biomatik) and two leaves per sample were snap frozen in liquid nitrogen. In barley, 10 leaf discs of 0.5 mm diameter per well were harvested from the tip of the third leaf of two weeks old plants and washed in H_2_O for 30 minutes in a 24-well plate. Leaf discs were then elicited with mock (water) or 1 µM flg22 and samples were collected and snap frozen. RNA was extracted using TRIzol™ (Invitrogen™) according to the manufacturer’s protocol. ReliaPrep™ RNA Miniprep Systems were used for RNA extraction for RNA Illumina sequencing according to the manufacturer’s instructions. After DNase I treatment (ThermoFisher), 1 µg of total RNA was used for cDNA synthesis using the RevertAid H Minus cDNA Synthesis Kit (ThermoFisher). Real-time quantitative PCR (RT-qPCR) was performed on cDNA samples using iQ SYBR Green Supermix (Bio-Rad) with primers listed in Supplementary Table S8 on a CFX Connect Real-Time PCR Detection System (Bio-Rad). The obtained Cq values were normalized to the Cq values of the housekeeping genes *AtUBIQUITIN 5* and *HvUBIQUITIN* relative to the WT mock treatment.

### ROS burst assays in Arabidopsis and barley

For ROS assays, 5 mm diameter leaf discs were obtained from 4-5-week-old Arabidopsis leaves and the upper part of the second leaves of 14-day-old barley plants. After washing with ddH_2_O, the discs were placed in 96-well plates and incubated overnight at 22 °C under aluminum foil. For Arabidopsis, the water was replaced with a solution containing 20 µM L-012, 2 µg ml^-1^ peroxidase (HRP), and 0.5 µM flg22 according to (Felix et al., 1999). In barley, leaf discs were first incubated for 30 min with the solution containing 10 µM L-012, 2 µg ml^-1^ HRP, and then 100 nM flg22 was added to the solution, as per (Scheler et al., 2016). The elicitor was not added in mock controls. Luminescence was measured using a microplate luminometer (Centro, LB960).

### Whole genome sequencing and off-target variant detection

High molecular weight (HMW) genomic DNA was isolated from four week old Arabidopsis WT, *tir-only-d1*, and *tir-only-d2* plants using 1.5 g of flash frozen leaf tissue with the NucleoBond HMW DNA kit (Macherey Nagel). Then DNA was quantifed (Qubit HS, Thermo), the size was assessed by capillary electrophoresis (Agilent FEMTOpulse) and 3 µg were then fragmented with Megaruptor 3 (Diagenode). PacBio libraries were prepared with an SMRTbell prep kit 3.0 according to the recommendations of the vendor (Pacific Biosciences). Then libraries were additionally enriched for larger library fragments (BluePippin, Sage Science). Trinary complexes (library-sequencing primer-Revio SPRQ polymerase) were prepared and sequenced with Revio SPRQ chemistry as a pool on a single Revio SMRT cell with 25 mio. ZMWs for 30h. Next, HiFi data was generated on the sequencer directly after the sequencing process. The Pacbio hifi long reads from Col-0 wildtype and the two mutant lines (d1, d2) were aligned to *A. thaliana* genome TAIR10 (Lamesch et al., 2012) using Minimap2 (v 2.28-r1209) (Li, 2018) and converted to BAM files using Samtools (v 1.16.1) (Danecek et al., 2021), with default parameters. With the BAM file as input, the variants were detected by DeepVariant (v 1.9.0) (Poplin et al., 2018) with default parameters. The variants called in the mutant lines were filtered out by the following steps: 1. also called in the wild type using BEDtools (v2.30.0) (Quinlan and Hall, 2010), 2. that are not marked as “PASS” by DeepVariant, 3. that do not have genotype quality (GQ) > 20, depth (DP) > 90 and variant allele frequency > 0.9 (VAF), 4. variants that occur in repeat regions. Steps 2 to 4 were carried out by custom python scripts. The effects of the variants were identified using SnpEff (Cingolani et al., 2012). The variants were manually confirmed by the Integrative Genomics Viewer (v2.19.4) (Robinson et al., 2011).

All the scripts used for this process can be found in: https://github.com/LeeeTak/variantcalling_DeepVariant_Laessle_etal

### Transcriptomic sequencing and analyses

For transcriptome analysis of barley Golden Promise and mutants (*eds1-pad4-d1*, *eds1-pad4-d2*, *tir-only-1*, *tir-only-2*), 20 million reads per sample were generated using Illumina technology. FastQC was used for quality control, and Trimmomatic trimmed the reads with parameters: leading:3, trailing:3, sliding window:4:20, minlen:36. The trimmed reads were mapped to the barley genome (*Hordeum vulgare* Morex V3) using HISAT2, with the reference from Ensembl Genomes. HTSeq counted mapped reads at the gene level, and DESeq2 in R performed differential expression analysis. Principal component analysis (PCA) was conducted using plotPCA() function from DESeq2 on the top 500 genes. The analysis focused on contrasts mock vs flg22 treatment for each genotype. Genes with an adjusted p-value < 0.05 and log_2_ fold change > 1 were considered significantly differentially expressed.

GO term enrichment analysis was performed using a hypergeometric test in R, with gene annotations from (Mascher et al., 2021) for barley using Arabidopsis, Chlamydomonas, and rice or NCBI blast. The hypergeometric test calculated the probability of observing at least k genes with a specific GO term in a sample of size n from a population of size N with m genes having the GO term. The phyper function in R implemented the test, and p-values were adjusted for multiple testing using the Benjamini-Hochberg method to control the false discovery rate (FDR).

### Generation of Arabidopsis and barley CRISPR/Cas9 mutant and Arabidopsis complementation lines

Guide RNA design was performed using CRISPR-P 2.0 (http://crispr.hzau.edu.cn/CRISPR2/) with *TIR*-*ONLY* CDS from Arabidopsis and barley and *EDS1*, *PAD4*, *ADR1* CDS from barley as input, with guide RNAs listed in Supplemental Table S8. Guide RNAs with high on-target scores were designed close to the start codon. The gRNA multiplex vectors were prepared as described in (Ordon et al., 2017; Ordon et al., 2020) and transformed into *Agrobacterium* GV3101 (pMP90) for Arabidopsis transformation and into *Agrobacterium* AGL1 strain for barley transformation. Arabidopsis Col-0 and *tir-only-d1* plants were transformed according to (Logemann et al., 2006). Barley Golden Promise WT was transformed according to (Tingay et al., 1997). Successful CRISPR mutagenesis was validated by PCR amplification (oligonucleotides listed in Supplemental Table S8) followed by Sanger sequencing. Homozygous barley *eds1 pad4* double and *adr1* single mutant lines (*ep-d1* and *ep-d2*; *adr1-1* and *adr1-2*) and Arabidopsis *tir-only* mutant were selected from which the Cas9 construct had been removed from the genome. Complementation was validated by gene expression and Western blot analysis.

### Phase separation microscopy

To study phase separation, CDS of *HvTIR*, *HvTIR^bb^*, *AtTIR*, and *AtTIR^bb^* were inserted into the pCHF3-GFP vector via KpnI and SalI digestion (Song et al., 2024). Microscopy of the transiently expressed constructs in *N. benthamiana* was performed following the method described by (Song et al., 2024).

### Statistics

Statistical analyses were performed using R software. Normality of residuals was assessed with the Shapiro-Wilk test, and homogeneity of variance was assessed with the Fligner-Killeen test. When assumptions were met, ANOVA was performed with aov(), followed by Tukey’s HSD post hoc test with TukeyHSD(). If normality was not achieved, data were log-transformed. For non-normal or heterogeneous data, the Kruskal-Wallis test was used with kruskal.test(), with pairwise comparisons using the Nemenyi post hoc test from the PMCMRplus package. A small random value was added to each measurement to avoid ties. For pairwise comparisons, a t-test was used for normal data, otherwise the Wilcoxon signed-rank test was used.

## Acknowledgements

We thank Chunpeng An (MPIPZ, Cologne) for kindly providing *Bgh* K1 primers and Fantin Mesny (University of Cologne, Cologne) for advice on RNA seq analysis. We thank Paul Schulze-Lefert (MPIPZ, Cologne) for valuable discussions and insightful comments on the manuscript. This study was supported by the Max Planck Society. H.L. and O.J. acknowledge International Max Planck Research School (IMPRS) support. J.E.P. and F.L. acknowledge support from iHEAD (NRW Profilbildung ID: PB22-025A). J.E.P., H.L. and F.L were further supported by Deutsche Forschungsgemeinschaft (DFG) CRC 1403 - project number 414786233 and Sino-German Mobility grant M-0275. L.L. was supported by the National Natural Science Foundation of China (grant no. 32470301). L.W. was supported by the National Natural Science Foundation of China (grant nos. 32525013 and 32270304).

## Author contributions

H.L., F.L. and J.E.P. designed the research; H.L., O.J., J.C., S.H., H.L., Y.C., J.B., J.J. performed experiments; H.L., L.L., W.S., J.J., B.H., T.L., L.W. analyzed the data; H.L. prepared figures and final datasets; H.L., F.L. and J.E.P wrote the manuscript with contributions from all authors.

## Supplementary Data

Supplementary Table S1. Whole genome sequencing identifies high-confidence variants in Arabidopsis *tir-only-d1* and *tir-only-d2* compared to the reference Col-0

Supplementary Table S2. Flg22-responsive genes (75) significantly upregulated in barley WT but not in the *eds1-pad4* and *tir-only* mutants

Supplementary Table S3. GO term enrichment for 75 flg22-responsive genes upregulated in barley WT but not in *eds1-pad4* and *tir-only* mutants

Supplementary Table S4. Flg22-responsive genes (25) significantly upregulated in barley in *eds1-pad4* and *tir-only* mutants but not in WT

Supplementary Table S5. GO term enrichment for 25 flg22-responsive genes upregulated in barley in *eds1-pad4* and *tir-only* mutants but not in WT

Supplementary Table S6. Downregulated genes (20) in barley WT after flg22 treatment

Supplementary Table S7. Protein sequence alignment between *At*BON1 and predicted *Hv*BON

Supplementary Table S8. Oligo primers used in this study

## Data availability

Relevant gene identifiers are provided in the manuscript and supporting information. Raw RNA-seq and whole-genome sequencing data have been deposited in the NCBI Sequence Read Archive (SRA) under BioProject accession numbers PRJNA1469211 and PRJNA1469868, respectively.

