## Supplementary figures (1-9) and data (1) for "Conserved TIR-only proteins drive transcriptional defense and basal immunity in dicot and monocot plants"

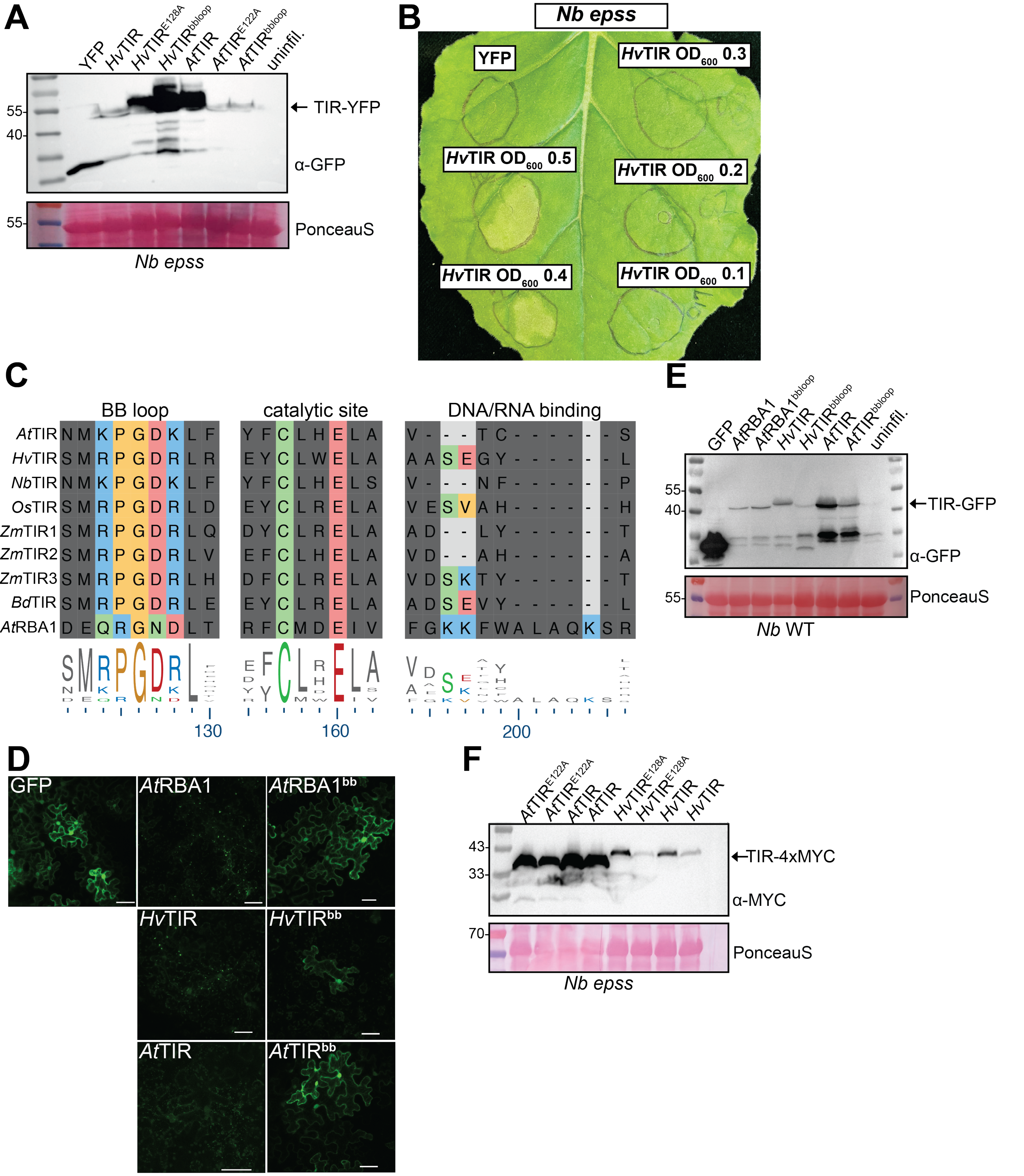
A
YFP
HvTIR
HvTIR E128A
HvTIR bbloop
AtTIR
AtTIR E122A
AtTIR bbloop
uninfil.
55-
40-
← TIR-YFP
α-GFP
55-
PonceauS
Nb epss
B
Nb epss
YFP
HvTIR OD600 0.3
HvTIR OD600 0.5
HvTIR OD600 0.2
HvTIR OD600 0.4
HvTIR OD600 0.1
C
BB loop
catalytic site
DNA/RNA binding
AtTIR
HvTIR
NbTIR
OsTIR
ZmTIR1
ZmTIR2
ZmTIR3
BdTIR
AtRBA1
130
160
200
D
GFP
AtRBA1
AtRBA1bb
HvTIR
HvTIRbb
AtTIR
AtTIRbb
E
GFP
AtRBA1
AtRBA1bbloop
HvTIR
HvTIRbbloop
AtTIR
AtTIRbbloop
uninfil.
55-
40-
←TIR-GFP
α-GFP
55-
PonceauS
Nb WT
F
AtTIR E122A
AtTIR E122A
AtTIR
AtTIR
HvTIR E128A
HvTIR E128A
HvTIR
HvTIR
43-
33-
←TIR-4xMYC
α-MYC
70-
PonceauS
Nb epss

**Supplementary Figure S1:** Cell death of conserved TIR-only in *Nicotiana benthamiana*. (**A**) Protein accumulation at 2 dpi in *Nb epss* from *Agrobacterium*-mediated expression of *At*TIR and *Hv*TIR, tagged with YFP and driven by the 35S promoter. Constructs include WT, non-catalytic (*At*TIR$^{E122A}$, *Hv*TIR$^{E128A}$), and BB-loop mutants (*At*TIR$^{bb}$, *Hv*TIR$^{bb}$) analyzed by Western blot with α-GFP antibodies. Ponceau S staining confirmed equal loading. Immunoblots repeated in four biological replicates. (**B**) Cell death in *Nb epss* induced by *Hv*TIR at varying $OD_{600}$, observed at 3 dpi in three independent experiments. YFP served as negative control, infiltrated at an $OD_{600}$ of 0.5. (**C**) Sequence alignment of conserved TIR-only protein motifs from Arabidopsis (*At*TIR), barley (*Hv*TIR), *N. benthamiana* (*Nb*TIR), rice (*Os*TIR), *Zea mays* (*Zm*TIR1-3), *Brachypodium distachyon* (*Bd*TIR) and Arabidopsis *At*RBA1 using Clustal Omega. Key amino acids highlighted. (**D**) Subcellular localization of *Hv*TIR, *Hv*TIR$^{bb}$, *At*TIR, *At*TIR$^{bb}$, *35S::AtRBA1-GFP* (*At*RBA1, as a positive control), *35S::AtRBA1$^{bb}$* (*At*RBA1$^{bb}$, as a negative control) and *35S::GFP* (GFP, as a negative control) in 4-5-weeks-old *Nb epss* leaves at 2 days after infiltration of *Agrobacteria* expressing the indicated constructs. Confocal images are representative of three independent experiments (Ten leaves from five plants were detected for each infiltration). Scale bars = 50 μm. (**E**) Protein accumulation of C-terminal GFP-tagged constructs used in the confocal microscopy. *Agrobacterium*-mediated expression of *35S::GFP* (control), *35S::AtRBA1-GFP*, *35S::AtRBA1*$^{bbloop}$-GFP, *35S::HvTIR-GFP*, *35S::HvTIR*$^{bbloop}$-GFP, *35S::AtTIR-GFP*, and *35S::AtTIR*$^{bbloop}$-GFP, alongside an uninfiltrated control in *N. benthamiana* WT. Proteins were detected by Western blot using α-GFP antibodies; Ponceau S staining served as a loading control. The immunoblot was repeated once. (**F**) Protein accumulation of C terminal 4xMYC tagged constructs used for small molecule measurements in *N. benthamiana epss*. Constructs (*35S::HvTIR-4xMYC*, *35S::HvTIR*$^{E128A}$*-4xMYC*, *35S::AtTIR-4xMYC*, and *35S::AtTIR*$^{E122A}$*-4xMYC*) were transiently expressed and detected by Western blot using α-MYC antibodies; Ponceau S staining served as loading control. The immunoblot was repeated once.

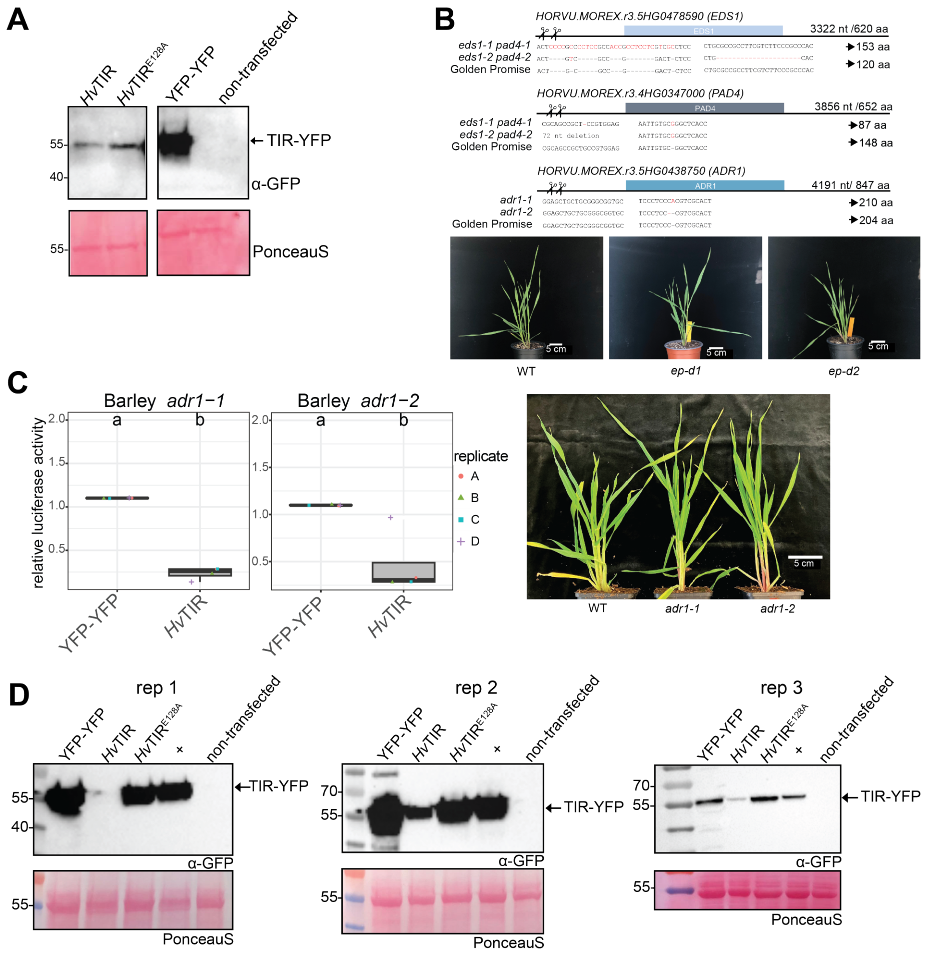

A
HvTIR
HvTIR E128A
YFP-YFP
non-transfected
TIR-YFP
α-GFP
PonceauS
B
HORVU.MOREX.r3.5HG0478590 (EDS1)
3322 nt /620 aa
eds1-1 pad4-1
eds1-2 pad4-2
Golden Promise
153 aa
120 aa
HORVU.MOREX.r3.4HG0347000 (PAD4)
3856 nt /652 aa
87 aa
148 aa
HORVU.MOREX.r3.5HG0438750 (ADR1)
4191 nt/ 847 aa
adr1-1
adr1-2
210 aa
204 aa
5 cm
WT
ep-d1
ep-d2
C
Barley adr1−1
Barley adr1−2
relative luciferase activity
replicate
A
B
C
D
YFP-YFP
HvTIR
WT
adr1-1
adr1-2
D
rep 1
rep 2
rep 3
YFP-YFP
HvTIR
HvTIR E128A
non-transfected
TIR-YFP
α-GFP
PonceauS

**Supplementary Figure S2:** Barley *eds1-pad4* and *adr1* mutants generated in this study. (**A**) Western blot of barley protoplast extracts transfected with pUBQ:luciferase and pUbi vectors containing *Hv*TIR and its inactive form *Hv*TIR$^{E128A}$, all tagged with YFP. Blots at 16 h post-transfection using α-GFP antibodies. Protein ladder on the left; expected sizes on the right. YFP and non-transfected controls included. Ponceau S staining confirmed equal loading. Immunoblot repeated once. (**B**) CRISPR/Cas9 targeting of *EDS1* (HORVU.MOREX.r3.5HG0478590), *PAD4* (HORVU.MOREX.r3.4HG0347000), and *ADR1* (HORVU.MOREX.r3.5HG0438750) in barley. *EDS1*, *PAD4* and *ADR1* were each targeted using two different guide RNAs (gRNAs) Black lines represent gene sequences with genomic nucleotide and protein lengths indicated. Scissors indicate gRNA target sites. Aligned sequences show DNA mutations and resulting protein sizes in amino acids. Photographs of 4-week-old WT, *eds1-pad4* double mutants, and *adr1* single mutants grown in greenhouse conditions. (**C**) Cell death assay in 7-9 days old barley *adr1-1* and *adr1-2* mutants transfected with pUBQ:luciferase and *Hv*TIR. Luciferase activity, measured 16 h post-transfection, indicates cell viability, normalized to YFP controls. Experiment repeated four times; dots represent technical replicates. Constructs with different letters above boxplots show significant differences (Kruskal-Wallis test with Nemenyi post-hoc, $\alpha = 0.05$, $n = 4$). (**D**) Expression of YFP-tagged *Hv*TIR and *Hv*TIR$^{E128A}$ in barley (WT) protoplasts for small molecule detection. Golden Promise WT protoplasts were transfected, and Western blot analysis was performed 16 h post-transfection using α-GFP antibodies. Protein ladder on the left; expected sizes on the right. YFP and the conserved TIR-only from rice (used here solely as positive Western blot control (+); (Wu et al., 2024)) and non-transfected controls included. Ponceau S staining confirmed equal loading. Experiment repeated three times.

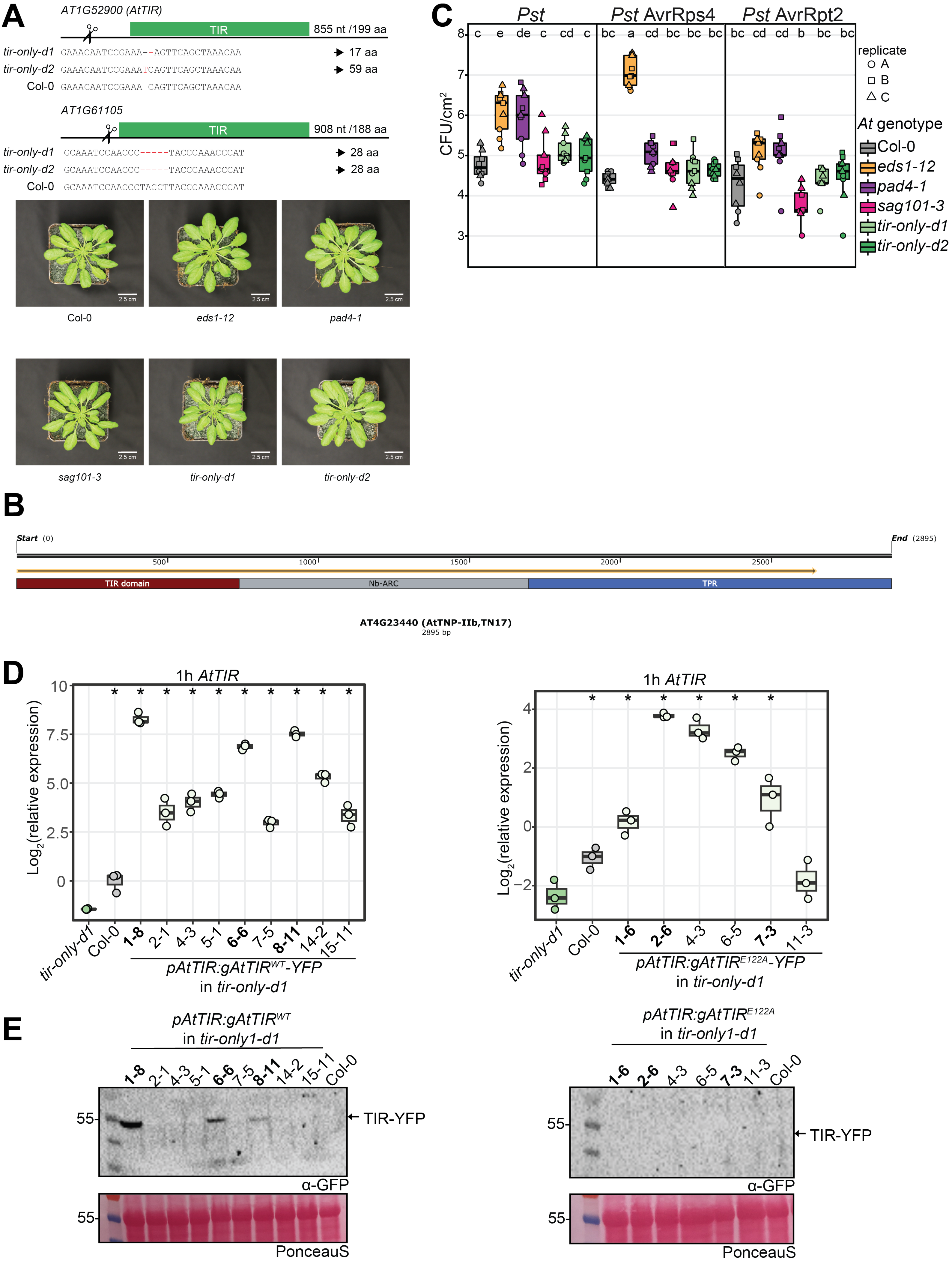
A
AT1G52900 (AtTIR)
TIR
855 nt /199 aa
tir-only-d1 GAAACAATCCGAAA--AGTTCAGCTAAACAA 17 aa
tir-only-d2 GAAACAATCCGAAATCAGTTCAGCTAAACAA 59 aa
Col-0 GAAACAATCCGAAA-CAGTTCAGCTAAACAA
AT1G61105
TIR
908 nt /188 aa
tir-only-d1 GCAAATCCAACCC-----TACCCAAACCCAT 28 aa
tir-only-d2 GCAAATCCAACCC-----TACCCAAACCCAT 28 aa
Col-0 GCAAATCCAACCCTACCTTACCCAAACCCAT
2.5 cm
Col-0
eds1-12
pad4-1
sag101-3
tir-only-d1
tir-only-d2
B
Start (0)
End (2895)
500
1000
1500
2000
2500
TIR domain
Nb-ARC
TPR
AT4G23440 (AtTNP-IIb,TN17)
2895 bp
C
Pst
Pst AvrRps4
Pst AvrRpt2
c e de c cd c
bc a cd bc bc bc
bc cd cd b bc bc
CFU/cm²
replicate
A
B
C
At genotype
Col-0
eds1-12
pad4-1
sag101-3
tir-only-d1
tir-only-d2
D
1h AtTIR
Log2(relative expression)
tir-only-d1
Col-0
1-8
2-1
4-3
5-1
6-6
7-5
8-11
14-2
15-11
pAtTIR:gAtTIRWT-YFP
in tir-only-d1
1h AtTIR
Log2(relative expression)
tir-only-d1
Col-0
1-6
2-6
4-3
6-5
7-3
11-3
pAtTIR:gAtTIRE122A-YFP
in tir-only-d1
E
pAtTIR:gAtTIRWT
in tir-only1-d1
1-8 2-1 4-3 5-1 6-6 7-5 8-11 14-2 15-11 Col-0
55
TIR-YFP
α-GFP
PonceauS
pAtTIR:gAtTIRE122A
in tir-only1-d1
1-6 2-6 4-3 6-5 7-3 11-3 Col-0
55
TIR-YFP
α-GFP
PonceauS

**Supplementary Figure S3:** Conserved TIR-only mutants and transgenic plants used in this study. (**A**,**B**) The two conserved *TIR-ONLY* genes in *Arabidopsis thaliana* (AT1G52900 (*AtTIR*) and AT1G61105) targeted by CRISPR/Cas9 are depicted with black lines, indicating their gene sequences along with genomic nucleotide and protein lengths. Scissors mark the gRNA target sites. Aligned sequences reveal DNA mutations and the corresponding protein sizes in amino acids. Photographs display 4-week-old Arabidopsis genotypes (Col-0, *eds1-12*, *pad4-1*, *sag101-3*, *tir-only-d1*, *tir-only-d2*) cultivated under greenhouse conditions. (**B**) Whole genome sequencing (WGS) identified a mutation in the C terminal TPR domain of AT4G23440 (*At*TNP IIb, TN17) in both Arabdiopsis conserved *tir-only* mutants. Domain annotation and sequence visualization were performed using InterPro and SNPAGene, respectively. The translated coding sequence of the mutant allele is indicated by a yellow arrow. (**C**) Growth of *Pseudomonas syringae* pv. *tomato* (*Pst*) DC3000 in 4-5 weeks old *Arabidopsis thaliana* Col-0, *eds1-12*, *pad4-1*, *sag101-3*, and *tir-only* mutants (*tir-only-d1*, *tir-only-d2*). Leaves were syringe-infiltrated with the *Pst* strain and *Pst* strains carrying AvrRps4, and AvrRpt2 ($OD_{600}$=0.0005). Bacterial titers were measured three days post-infection, presented as CFU/cm² in boxplots. Colors denote genotypes, with individual measurements as shapes. Different letters in boxplot indicate significant differences (Tukey HSD, $\alpha = 0.01$, $n = 9$). (**D**) *AtTIR* expression was measured in four-week-old Arabidopsis Col-0, *tir-only-d1*, and selected complementation lines one hour after 0.5 µM flg22 treatment by RT-qPCR. Results were normalized to mock-treated Col-0 with asterisks indicating statistically significance difference compared to mock-treated *tir-only-d1* (Wilcoxon signed-rank test, $p < 0.05$, $n = 3$). (**E**) Western blot analysis of the same samples using α-GFP antibody confirmed protein expression. Col-0 served as a negative control, with protein levels indicated and equal loading verified by Ponceau S staining. (**D**,**E**) Analyses were repeated once, selected complementation lines are shown in bold.

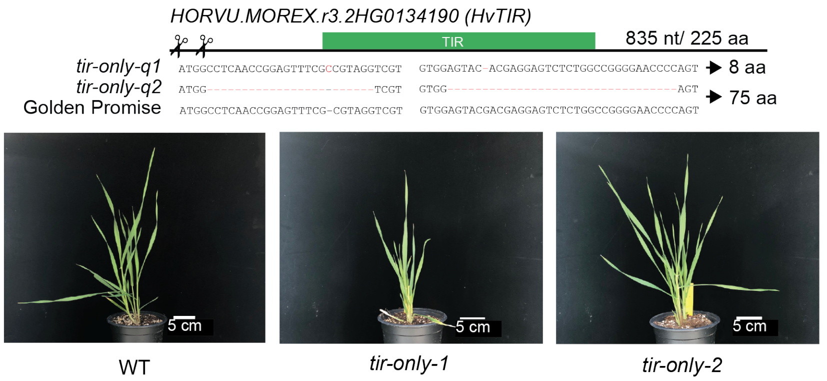


**Supplementary Figure S4:** Barley conserved TIR-only CRISPR mutant. The conserved *TIR-ONLY* gene in barley (HORVU.MOREX.r3.2HG0134190) targeted by CRISPR/Cas9 are depicted with black lines, indicating their gene sequences along with genomic nucleotide and protein lengths. Scissors mark the gRNA target sites. Aligned sequences reveal DNA mutations and the corresponding protein sizes in amino acids. Photographs display 4 weeks old barley genotypes (WT, *tir-only-1*, *tir-only-2*) cultivated under greenhouse conditions.

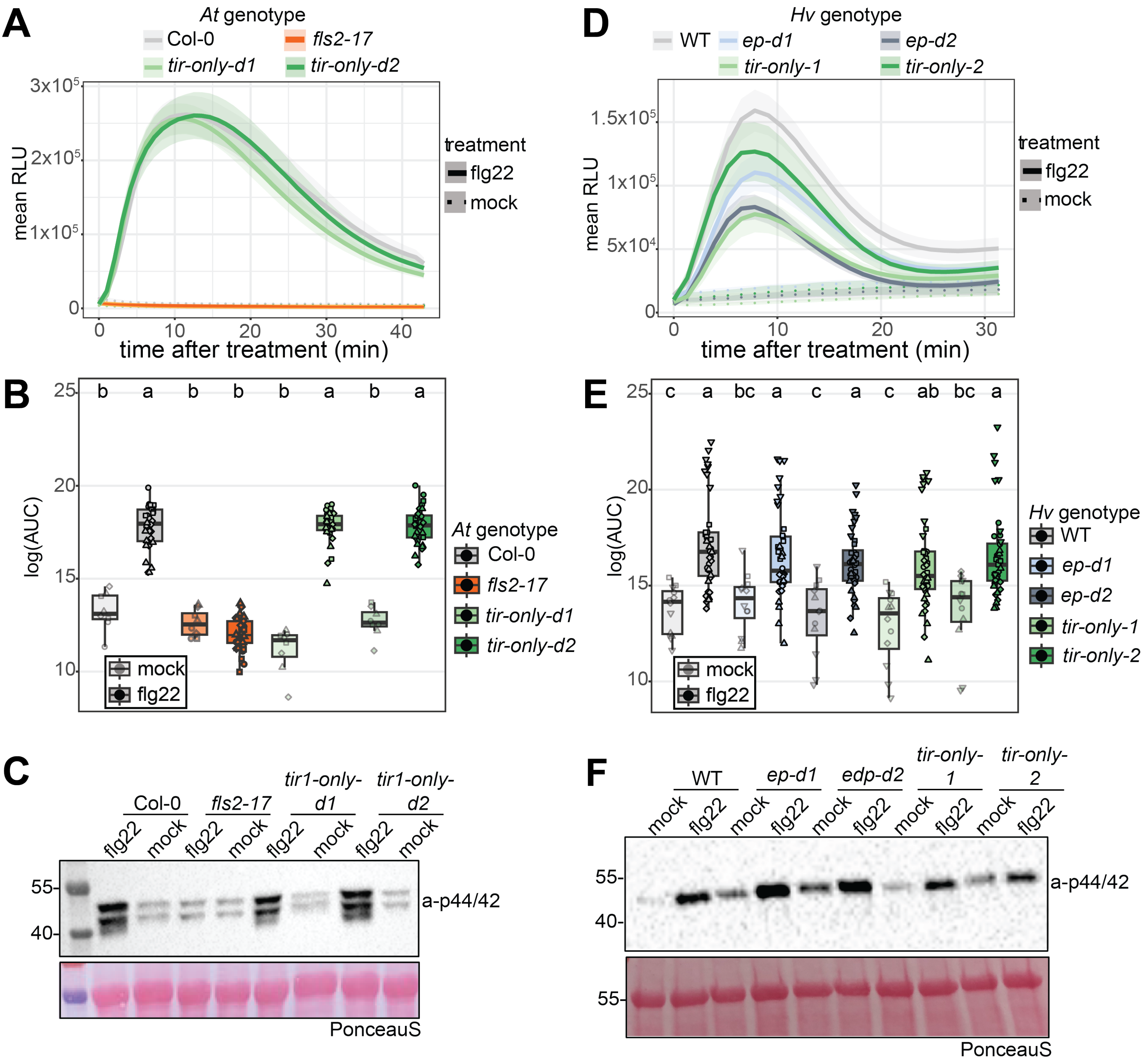

A
At genotype
Col-0
fls2-17
tir-only-d1
tir-only-d2
mean RLU
time after treatment (min)
treatment
flg22
mock
B
log(AUC)
b a b b b a b a
At genotype
Col-0
fls2-17
tir-only-d1
tir-only-d2
mock
flg22
C
Col-0
fls2-17
tir1-only-d1
tir1-only-d2
flg22 mock flg22 mock flg22 mock flg22 mock
55-
40-
a-p44/42
PonceauS
D
Hv genotype
WT
ep-d1
ep-d2
tir-only-1
tir-only-2
mean RLU
time after treatment (min)
treatment
flg22
mock
E
log(AUC)
c a bc a c a c ab bc a
Hv genotype
WT
ep-d1
ep-d2
tir-only-1
tir-only-2
mock
flg22
F
WT
ep-d1
edp-d2
tir-only-1
tir-only-2
mock flg22 mock flg22 mock flg22 mock flg22 mock flg22
55-
40-
a-p44/42
55-
PonceauS

**Supplementary Figure S5:** *At*TIR and *Hv*TIR do not contribute to flg22-induced ROS production and MAPK activation in Arabidopsis and barley. (**A,D**) ROS production measured as relative light units (RLUs) in 4 weeks old *Arabidopsis thaliana* (Col-0, *fls2-17*, *tir-only-d1*, *tir-only-d2*) and 2 weeks old barley (WT, *ep-d1*, *ep-d2*, *tir-only-1*, *tir-only-2*) following flg22 treatment (0.5 µM in Arabidopsis; 1 µM in barley). Solid lines represent flg22 treatment; dashed lines indicate mock treatment. Colors denote genotypes. Experiments repeated four times for Arabidopsis and five times for barley. (**B,E**) Boxplots of log-transformed area under the curve (AUC) for 30-min ROS measurements. Genotypes with different letters above boxplots show significant differences (Kruskal-Wallis post-hoc Nemenyi test, $p = 0.05$). Sample sizes: $n = 32$ for flg22 in Arabidopsis, $n = 40$ for flg22 in barley, $n = 8$ for mock in Arabidopsis, and $n = 10$ for mock in barley. (**C,F**) MAPK activation in 4 weeks old Arabidopsis and 2 weeks old barley genotypes assessed by phospho-specific p44/42 antibody blotting, 15 min post-flg22 treatment (0.5 µM in Arabidopsis; 1 µM in barley). Protein ladder on the left. Equal loading confirmed by Pierce assay and Ponceau S staining. Immunoblots repeated three times.

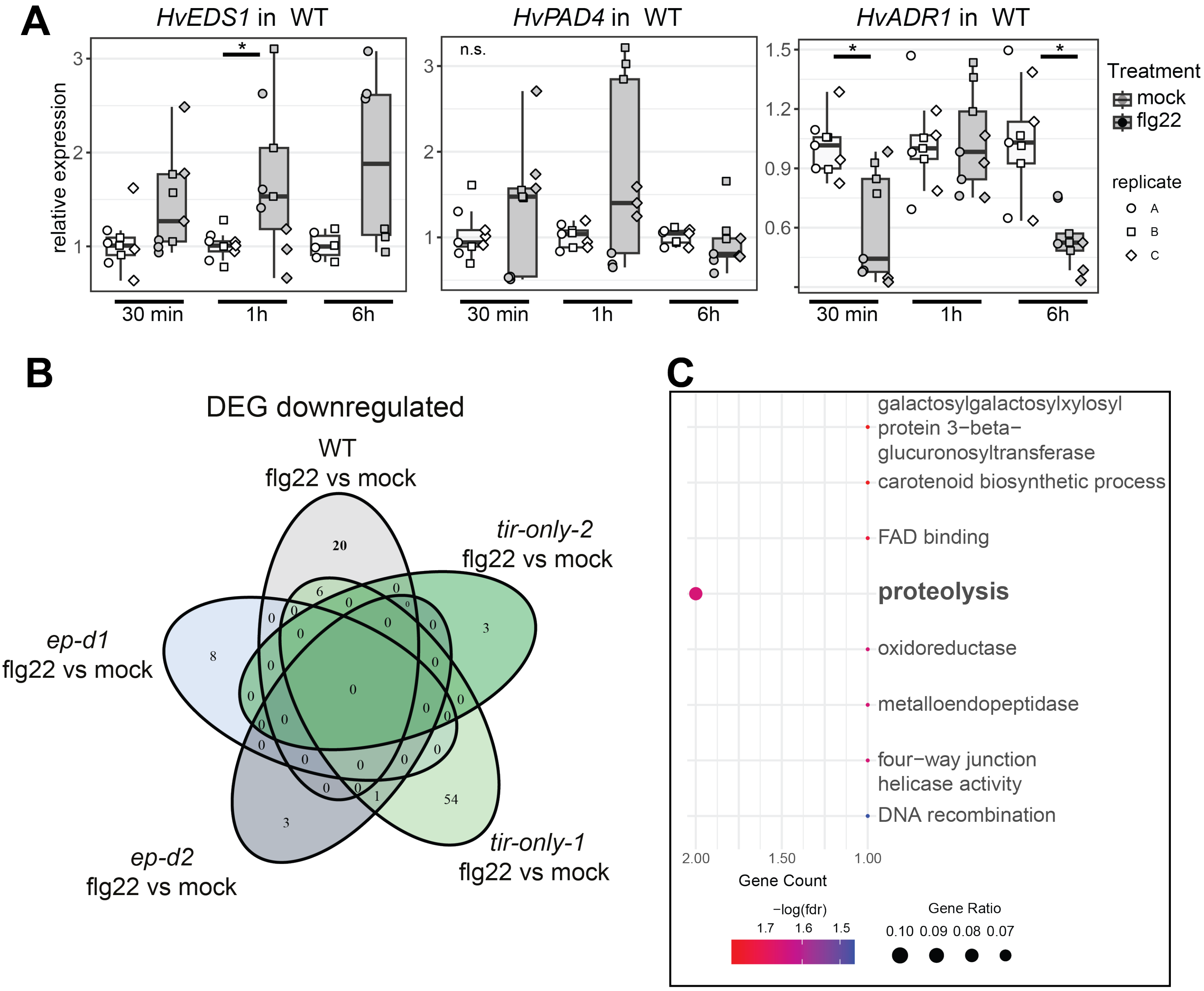


**Supplementary Figure S6:** Transcriptome changes 1h after flg22 treatment in barley conserved *eds1-pad4* and *tir-only* mutants. (**A**) RT-qPCR analysis of *HvEDS1*, *HvPAD4*, and *HvADR1* expression in two-week-old WT barley following 1 µM flg22 treatment at 30 min, 1 h, and 6 h. The experiment was conducted independently three times. Relative quantification (RQ) was calculated relative to mock-treated samples at each time point and normalized to *UBIQUITIN* expression. Asterisks indicate significant expression differences between flg22 and mock treatments (Wilcoxon signed-rank test, $p < 0.05$, $n = 9$). (**B**) Differential gene expression (DEG) analysis was performed for barley WT, *ep-d1*, *ep*-d2, *tir-only-1*, and *tir-only-2* genotypes, comparing flg22 treatment with mock treatment for each genotype. Genes were considered significant if they had an adjusted p-value < 0.05 and a log2 fold-change > 1. The Venn diagram, generated using the VennDiagram package, illustrates the overlap of downregulated genes between genotypes. (**C**) GO term analysis was performed on the significantly downregulated genes in WT using hypergeometric testing. Significance is represented by different colored dots (FDR < 0.05), and the size of the dots indicates the gene ratio. Gene annotations of twenty genes can be found in Supplementary table S6.

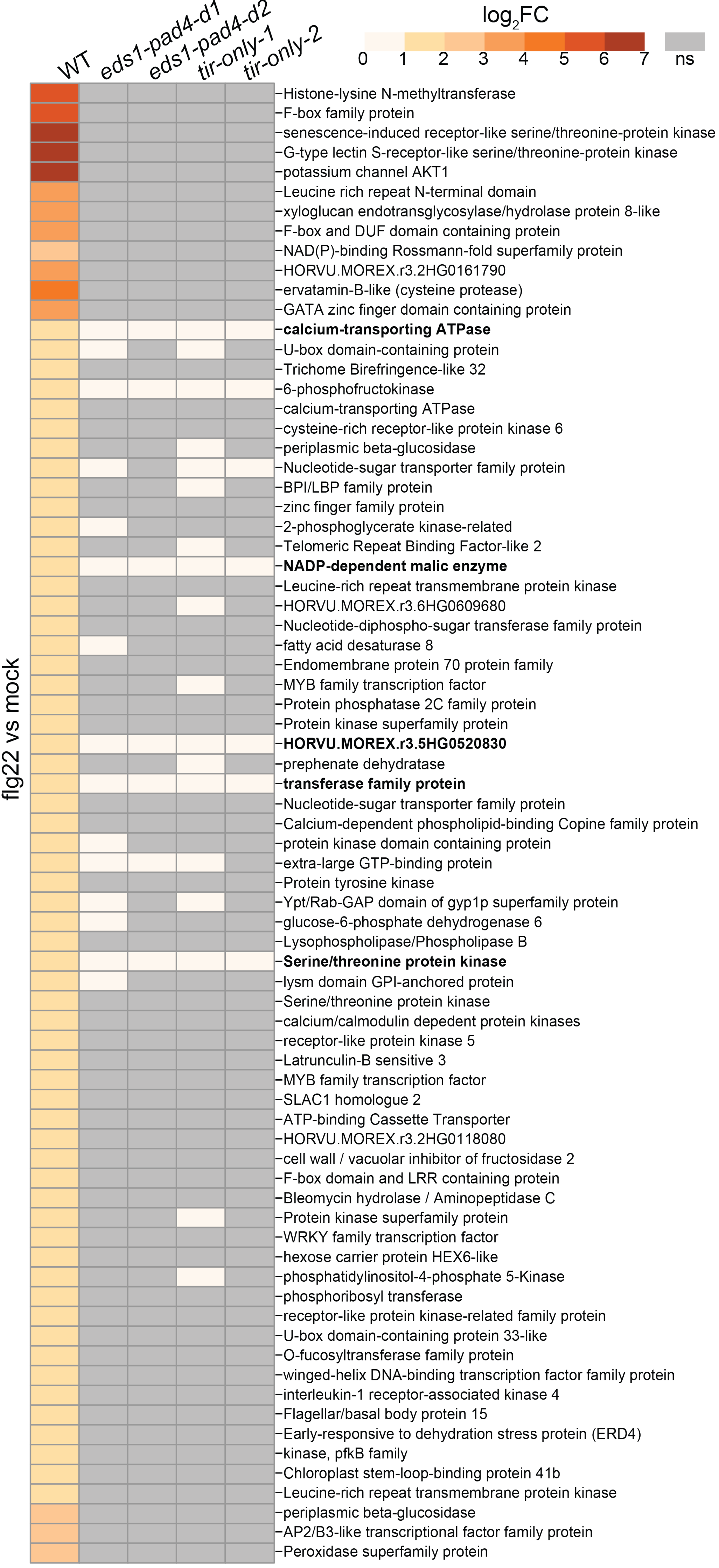
WT
eds1-pad4-d1
eds1-pad4-d2
tir-only-1
tir-only-2
log2FC
0
1
2
3
4
5
6
7
ns
flg22 vs mock
Histone-lysine N-methyltransferase
F-box family protein
senescence-induced receptor-like serine/threonine-protein kinase
G-type lectin S-receptor-like serine/threonine-protein kinase
potassium channel AKT1
Leucine rich repeat N-terminal domain
xyloglucan endotransglycosylase/hydrolase protein 8-like
F-box and DUF domain containing protein
NAD(P)-binding Rossmann-fold superfamily protein
HORVU.MOREX.r3.2HG0161790
ervatamin-B-like (cysteine protease)
GATA zinc finger domain containing protein
calcium-transporting ATPase
U-box domain-containing protein
Trichome Birefringence-like 32
6-phosphofructokinase
calcium-transporting ATPase
cysteine-rich receptor-like protein kinase 6
periplasmic beta-glucosidase
Nucleotide-sugar transporter family protein
BPI/LBP family protein
zinc finger family protein
2-phosphoglycerate kinase-related
Telomeric Repeat Binding Factor-like 2
NADP-dependent malic enzyme
Leucine-rich repeat transmembrane protein kinase
HORVU.MOREX.r3.6HG0609680
Nucleotide-diphospho-sugar transferase family protein
fatty acid desaturase 8
Endomembrane protein 70 protein family
MYB family transcription factor
Protein phosphatase 2C family protein
Protein kinase superfamily protein
HORVU.MOREX.r3.5HG0520830
prephenate dehydratase
transferase family protein
Nucleotide-sugar transporter family protein
Calcium-dependent phospholipid-binding Copine family protein
protein kinase domain containing protein
extra-large GTP-binding protein
Protein tyrosine kinase
Ypt/Rab-GAP domain of gyp1p superfamily protein
glucose-6-phosphate dehydrogenase 6
Lysophospholipase/Phospholipase B
Serine/threonine protein kinase
lysm domain GPI-anchored protein
Serine/threonine protein kinase
calcium/calmodulin depedent protein kinases
receptor-like protein kinase 5
Latrunculin-B sensitive 3
MYB family transcription factor
SLAC1 homologue 2
ATP-binding Cassette Transporter
HORVU.MOREX.r3.2HG0118080
cell wall / vacuolar inhibitor of fructosidase 2
F-box domain and LRR containing protein
Bleomycin hydrolase / Aminopeptidase C
Protein kinase superfamily protein
WRKY family transcription factor
hexose carrier protein HEX6-like
phosphatidylinositol-4-phosphate 5-Kinase
phosphoribosyl transferase
receptor-like protein kinase-related family protein
U-box domain-containing protein 33-like
O-fucosyltransferase family protein
winged-helix DNA-binding transcription factor family protein
interleukin-1 receptor-associated kinase 4
Flagellar/basal body protein 15
Early-responsive to dehydration stress protein (ERD4)
kinase, pfkB family
Chloroplast stem-loop-binding protein 41b
Leucine-rich repeat transmembrane protein kinase
periplasmic beta-glucosidase
AP2/B3-like transcriptional factor family protein
Peroxidase superfamily protein

**Supplementary Figure S7:** DEG upregulated in WT barley. Differential gene expression analysis was conducted in WT barley, *eds1-pad4*, and *tir-only* mutants comparing mock and flg22 treatments. Seventy-five genes with an adjusted p-value $< 0.05$ and $\log_2$ fold-change $> 1$ were identified as significantly upregulated in WT. The heatmap illustrates $\log_2$ fold-change expression levels across the genotypes, with non-significant changes shown in grey.

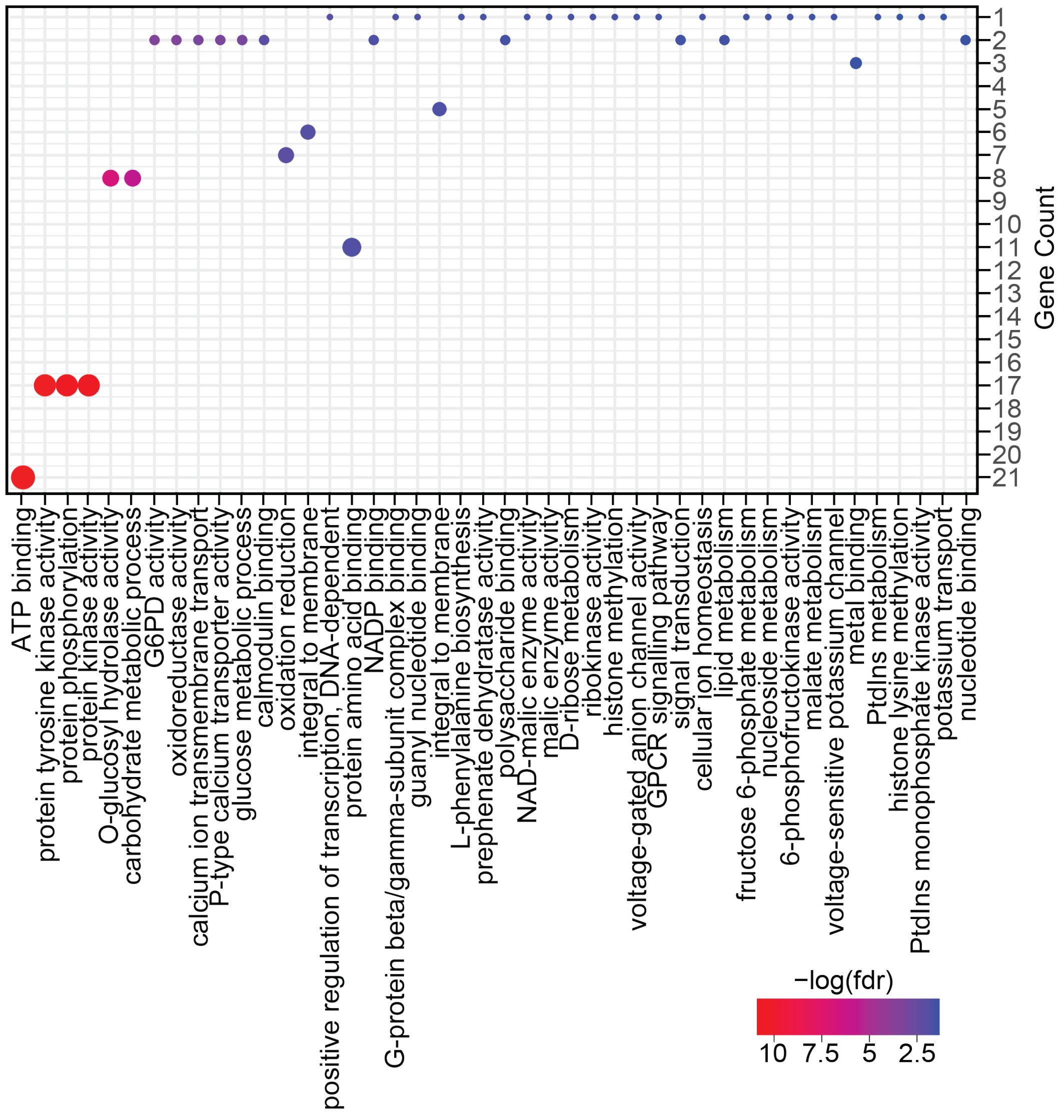


**Supplementary Figure S8:** GO term analysis of upregulated DEG in WT. GO term enrichment analysis of significantly upregulated genes in WT using hypergeometric testing. Dot color indicates significance (FDR < 0.05), and dot size reflects the gene ratio.

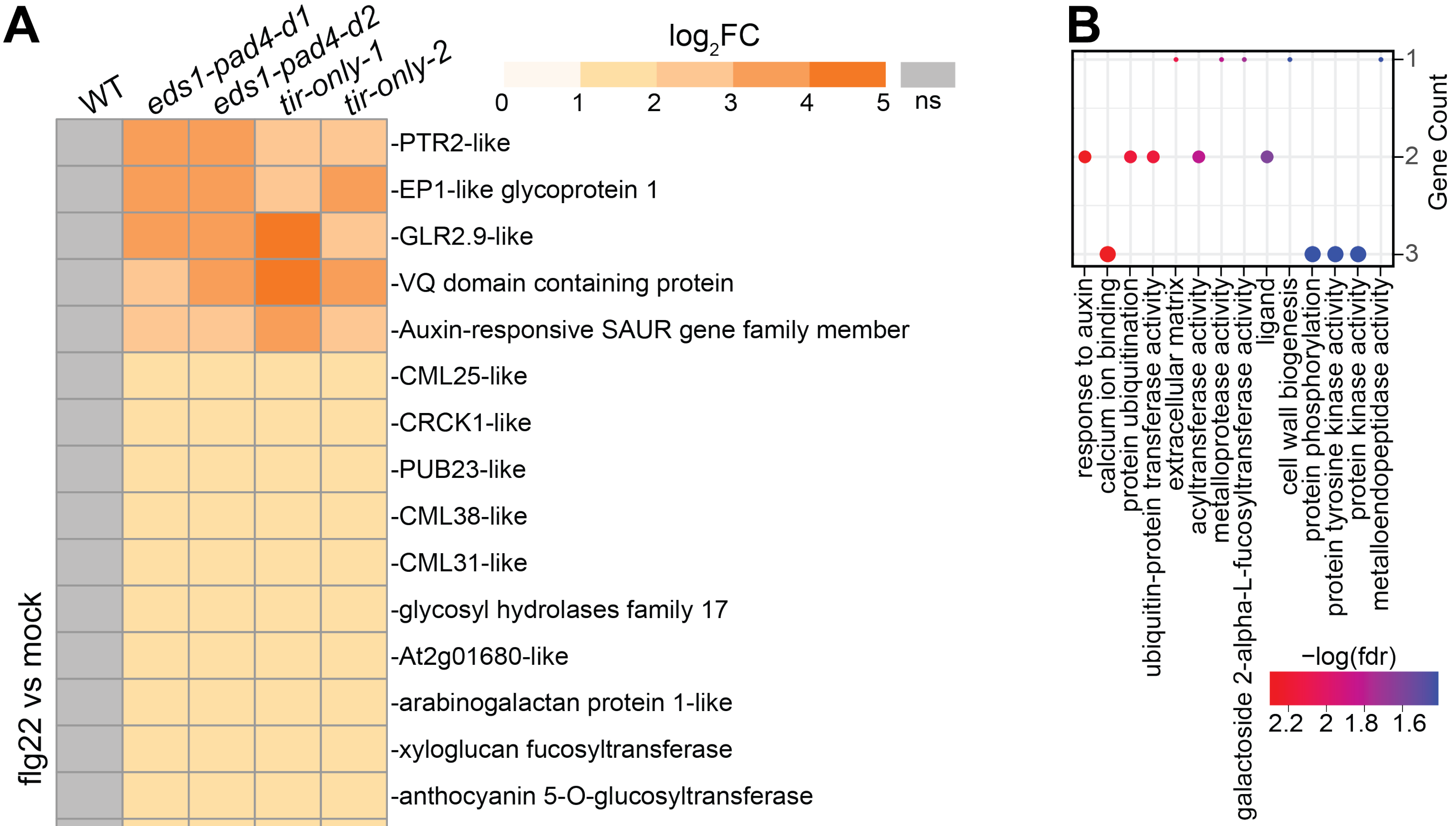


**Supplementary Figure S9:** Upregulated DEG in *eds1-pad4* and *tir-only* barley mutants. **(A)** Differential gene expression analysis was conducted in WT barley, *eds1-pad4*, and *tir-only* mutants comparing mock and flg22 treatments. Twenty-five genes with an adjusted p-value < 0.05 and $log_2$ fold-change > 1 were identified as significantly upregulated in *eds1-pad4* and *tir-only* mutants. The heatmap illustrates $log_2$ fold-change expression levels across the genotypes, with non-significant changes shown in grey. **(B)** GO term enrichment analysis of significantly upregulated genes *eds1-pad4* and *tir-only* mutants using hypergeometric testing. Dot color indicates significance (FDR < 0.05), and dot size reflects the gene ratio.

variants called with deepvariant
(https://www.nature.com/articles/nbt.4235)

Potential off-target mutation in
AT4G23440 (AtTNP-IIb, TN17)

Col0

d1

d2

variant cutoff:
GQ (genotype quality) > 20 (> 99% accuracy),
depth > 100,
variant allele frequency > 0.9

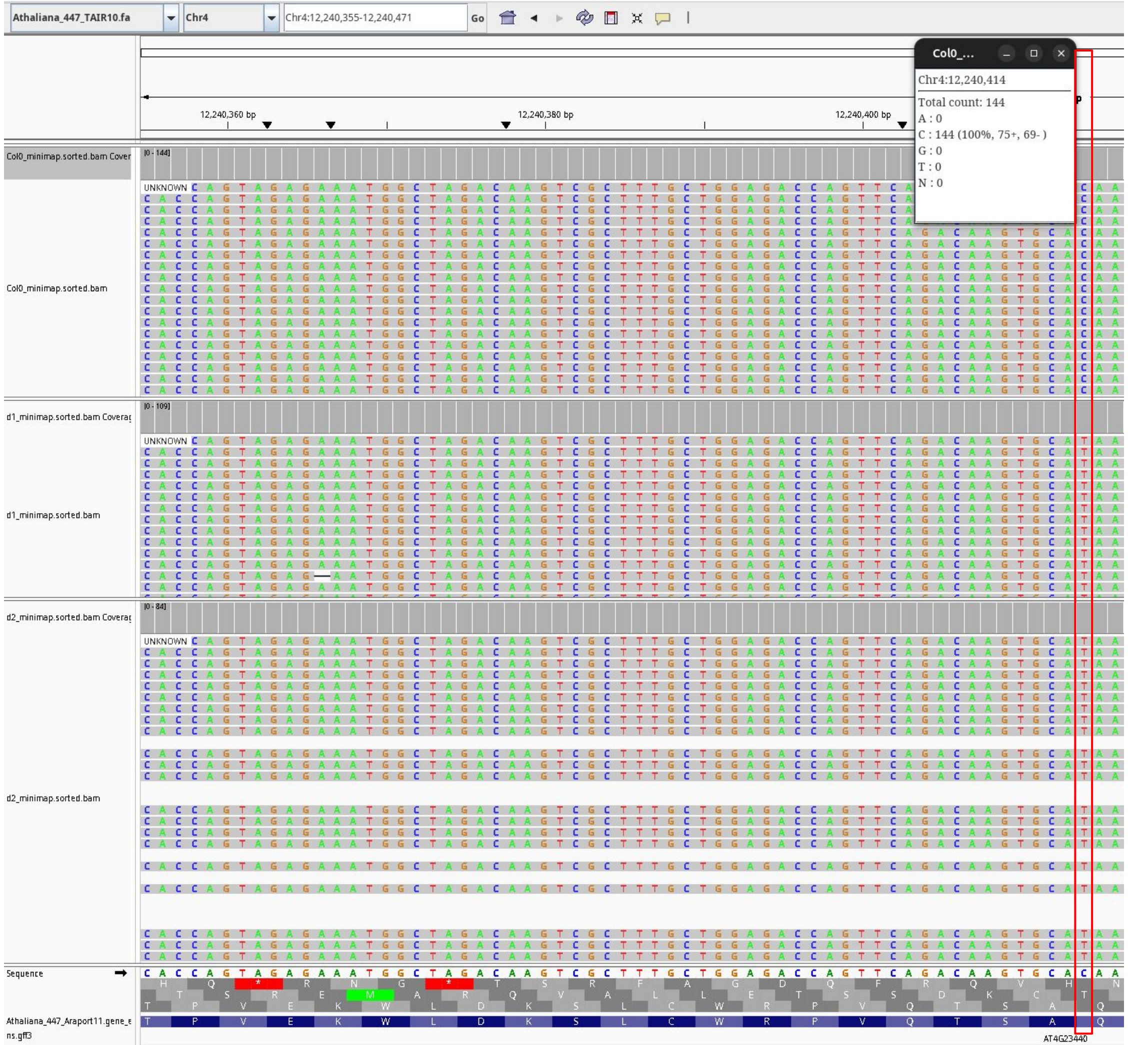


C > T causes
CAA (Q) to TAA (UAA,stop)

### Supplementary Data S1
